# Programming the lifestyles of engineered bacteria for cancer therapy

**DOI:** 10.1101/2022.07.15.500166

**Authors:** Shengwei Fu, Rongrong Zhang, Yanmei Gao, Jiarui Xiong, Ye Li, Lu Pu, Aiguo Xia, Fan Jin

## Abstract

Bacteria can be genetically engineered to act as therapeutic delivery vehicles in the treatment of tumors, killing cancer cells or activating the immune system. This is known as Bacteria-Mediated Cancer Therapy (BMCT). Tumor invasion, colonization and tumor regression are major biological events, which are directly associated with antitumor effects and are uncontrollable due to the influence of tumor microenvironments during the BMCT process. Here, we developed a genetic circuit for dynamically programming bacterial lifestyles (planktonic, biofilm or lysis), to precisely manipulate the process of bacterial adhesion, colonization and drug release in BMCT process, via hierarchical modulation of the lighting power density (LPD) of near-infrared (NIR) light. The deep tissue penetration of NIR offers us a modality for spatiotemporal and noninvasive control of bacterial genetic circuits in vivo. By combining computational modeling with high throughput characterization device, we optimized the genetic circuits in engineered bacteria to program the process of bacterial lifestyle transitions by altering the illumination scheme of NIR. Our results showed that programming intratumoral bacterial lifestyle transitions allows precise control of multiple key steps throughout the BMCT process, and therapeutic efficacy can be greatly improved by controlling the localization and dosage of therapeutic agents via optimizing the illumination scheme.

## INTRODUCTION

The hypoxia and immune-privileged tumor microenvironment is one of the biggest obstacles in the way of many cancer therapeutics[1–3], such as chemotherapy[4], radiotherapy[5], and immunotherapy[6, 7]. Nevertheless, this unique microenvironment of solid tumors is ideal for the colonization and proliferation of a variety of obligate anaerobes and facultative anaerobes[8–11], including *Listeria, Clostridium,* and *Salmonell*, which have been well studied and extensively used in the treatment of cancers. In addition to their inherent antitumor effects by stimulation of the innate immune system or secretion of natural antitumor products[12–15], bacteria were also engineered to act as therapeutic vehicles to deliver different payloads for improved in situ cancer therapy[16–22]. The engineered living therapeutics endowed with synthetic genetic circuits have shown advantages over conventional cancer therapies in terms of flexibility, specificity and predictability[23]. Bacteria-mediated cancer therapy (BMCT) has therefore emerged as a promising strategy for the treatment of cancers.

Tumor invasion, colonization and tumor regression are key biological events associated with antitumor effects in the process of BMCT[24]. Different approaches have been adopted to improve the therapeutic efficacy via manipulating these biological events, such as virulence attenuation[25], bacterial colonization enhancement[26, 27] and drug release strategies[28–30]. For example, ΔppGpp *S. typhimurium* strain, expressing tumor-specific ligands RGD peptide on the cell surface, improves bacterial tumor targeting and therapeutic efficacy[27]. Although these strategies resulted in improved therapeutic efficacy in mice, the situation is different when it comes to clinical study, which shows lower colonization densities and greater heterogeneity[31]. For most therapeutic agents like chemical drugs and cytotoxic proteins used in BMCT, therapeutic efficacy is dose-dependent, requiring higher colonization for better therapeutic efficacy. These clinical trials, on the other hand, suggest that intratumoral colonization stems from the bidirectional biological interactions between bacteria and the host tumor microenvironment (TME), which is dynamic and uncontrollable to some extent[32].

As tumor treatment is a long-term process that requires controlled and sustained release of therapeutic agents into the TME[28, 33], tumor regression is difficult to achieve by simply manipulating a certain biological event in BMCT, such as bacterial colonization and drug release. While the development of nanotechnology and the increased availability of versatile materials, including polymeric hydrogels and lipids, have opened up new possibilities for achieving sustained drug release[34], this remains a challenge in BMCT. Din and coworkers developed a synchronized lysis system that enables periodic drug production with the fluctuation in bacterial populations[28]. In this system, tumor growth is inhibited by the therapeutic agents released via bacterial lysis, while the process of bacterial colonization is uncontrollable and the density of bacterial colonization cannot exceed a threshold due to intrinsic lysis mechanism. Therefore, multiple injections are necessary to maintain the treatment for better therapeutic efficacy, but at the same time bring about pharmacological adverse effects.

Optogenetics allows for the control of cellular signaling in real time and is recently being employed for therapeutic applications[35]. Using optogenetics, we developed a genetic circuit in engineered bacteria that allows dynamic manipulation of bacterial lifestyles (planktonic, biofilm and lysis lifestyle) to precisely control the process of bacterial adhesion, colonization and drug release, with near-infrared (NIR) light in the BMCT process. The deep tissue penetration of NIR enables us a spatiotemporal and noninvasive control of genetic circuit[36, 37] and is widely used to trigger certain behaviors of bacteria in vivo[38, 39]. In addition, the LPD of NIR used to program the lifestyles of engineered bacteria H017 shows a reduction of 3 orders of magnitude compared to that of bacteria-based photothermal therapy (PTT)[40, 41], which enables widespread clinical use outside of dermatological indications. We also explored the potential of H017 for cancer therapy. Much less frequent injections of H017 can accomplish controllable drug release and tumor repression by dynamically programming the intratumoral bacterial lifestyle transitions via hierarchical modulation of the LPD of NIR. In all, the programmable lifestyle system enables multiple critical steps of the entire BMCT process to be precisely controlled, and expands the strategy for enhanced bacterial colonization and sustained drug release in solid tumors.

## RESULTS

### Attenuation of *P. aeruginosa* for cancer therapy

*Salmonella tyhimurium* and some other pathogens[42], which have been extensively studied for the treatment of tumors, have rarely been reported to colonize the lung. However, *Pseudomonas aeruginosa* can naturally colonize the lung in mouse models when inoculated intranasally (Table S1) and preferentially accumulate in solid tumors[43, 44] just like *Salmonella* and *E. coli* (Table S2). Besides, bacteriocins (exotoxin A, PE) produced by *P. aeruginosa* shows antitumor activities[45], and *P. aeruginosa* preparation (PAP) has already been used to inhibit the metastasis of lung cancer in clinical trials [46]. Taken together, we reasoned that *P. aeruginosa* has great potential for the treatment of primary lung cancer.

Attenuation of human pathogens is critical for BMCT. For *P. aeruginosa*, Vfr (virulence factor regulator) is a cAMP-binding transcriptional regulator that controls the production of multiple virulence factors on a global level[47]. *Vfr* mutant showed notably weakened cytotoxicity to host cells[48]. In addition, Type III secretion system (T3SS) exoenzyme S (ExoS) and exoenzyme T (ExoT) are involved in *P. aeruginosa* pathogenesis. *P. aeruginosa* mutant lacking both *exoS* and *exoT* has remarkably decreased ability to survive or spread systemically[49]. Based on previous studies, by triple deletion of *exoS*, *exoT* and *vfr* successively, we constructed a *P. aeruginosa* strain, which was significantly attenuated both in vitro and in vivo compared to the wild type strain (Fig. S1).

### Genetic circuit design for programming bacterial lifestyles

We designed three distinct lifestyles to implement different functions in BMCT process. The adhesion of bacteria to tissue surfaces is an essential step in bacterial invasions in BMCT. Thus, we speculated that setting the initial bacterial lifestyle to non-colonization (planktonic lifestyle) prior to invading tumor tissues can greatly reduce its toxicity to normal tissues. It is well known that the formation of biofilm prevents bacteria from neutrophil-mediated killing[50] and facilitates their adhesion to the surface of various tissues[51]. We hypothesized that intratumoral biofilm formation of *P. aeruginosa* may contribute to immune privilege, enhanced tumor colonization and improved therapeutic efficacy in BMCT. So, biofilm lifestyle was also designed. Extracellular polymeric substances (EPS) play a vital role in bacterial initial adhesion, surface colonization, biofilm formation and dispersal, and EPS production can be regulated by the bacterial secondary messenger c-di-GMP[52]. The intracellular c-di-GMP can be degraded by phosphodiesterase PA2133 and synthesized by BphS in a NIR-dependent manner[53]. Moreover, the LPD and illumination duration of NIR can be precisely manipulated in vitro. Therefore, with the genetic circuit consisting of a NIR sensor module and a c-di-GMP hydrolysis module, the bacterial intratumoral lifestyle can be tuned from planktonic (non-colonization) to biofilm (colonization) when irradiated with appropriate LPDs of NIR. To facilitate drug release after bacterial colonization of tumors, lysis lifestyle was designed as well. We firstly screened the lysis genes from different microorganisms (Fig. S2, Table S3) to obtain a lysis cassette *LKD* that can cause lysis of wild type *P. aeruginosa* PAO1 at a relatively low expression level, given that a low expression level of lysis gene can greatly reduce the burden of therapeutic bacteria and thus improve the stability of the inducible lysis system[54]. To avoid the leaky expression of the genes to induce the death of a small fraction of cells, the *LKD* was placed behind the terminator tR’ (Fig. S3), so that the *LKD* is activated by the production of anti-terminator protein Q which was driven by the c-di-GMP responsive promoter P*cdrA*. As the LPD increases, the production of Q protein reaches a threshold, leading to the lysis of bacteria and the release of antitumor drugs into TME. Besides, the expression noise of *Q* contributes to phenotypic variability and results in asynchronous lysis behavior. When the LPD of NIR is tuned to a lower level, the surviving bacteria continue to grow as seeds, allowing for the subsequent regulation of bacterial lifestyle transitions for sustainable drug synthesis and release.

By regulating the NIR-responsive c-di-GMP levels, the lifestyle-programmable cancer therapy system can be easily switched between distinct bacterial lifestyles (Fig. 1b). To be specific, under darkness or a low-LPD of NIR, bacteria are prevented from adhering to the tissue sites; when a medium-LPD of NIR is supplied, light-activated DGC (BphS) causes an increase in c-di-GMP level and subsequently enhances biofilm formation; upon illumination with a high-LPD of NIR, lysis is triggered, and the pore-forming anti-tumor toxin Hemolysin E (HlyE) is released to kill tumor cells.

**Figure 1.**
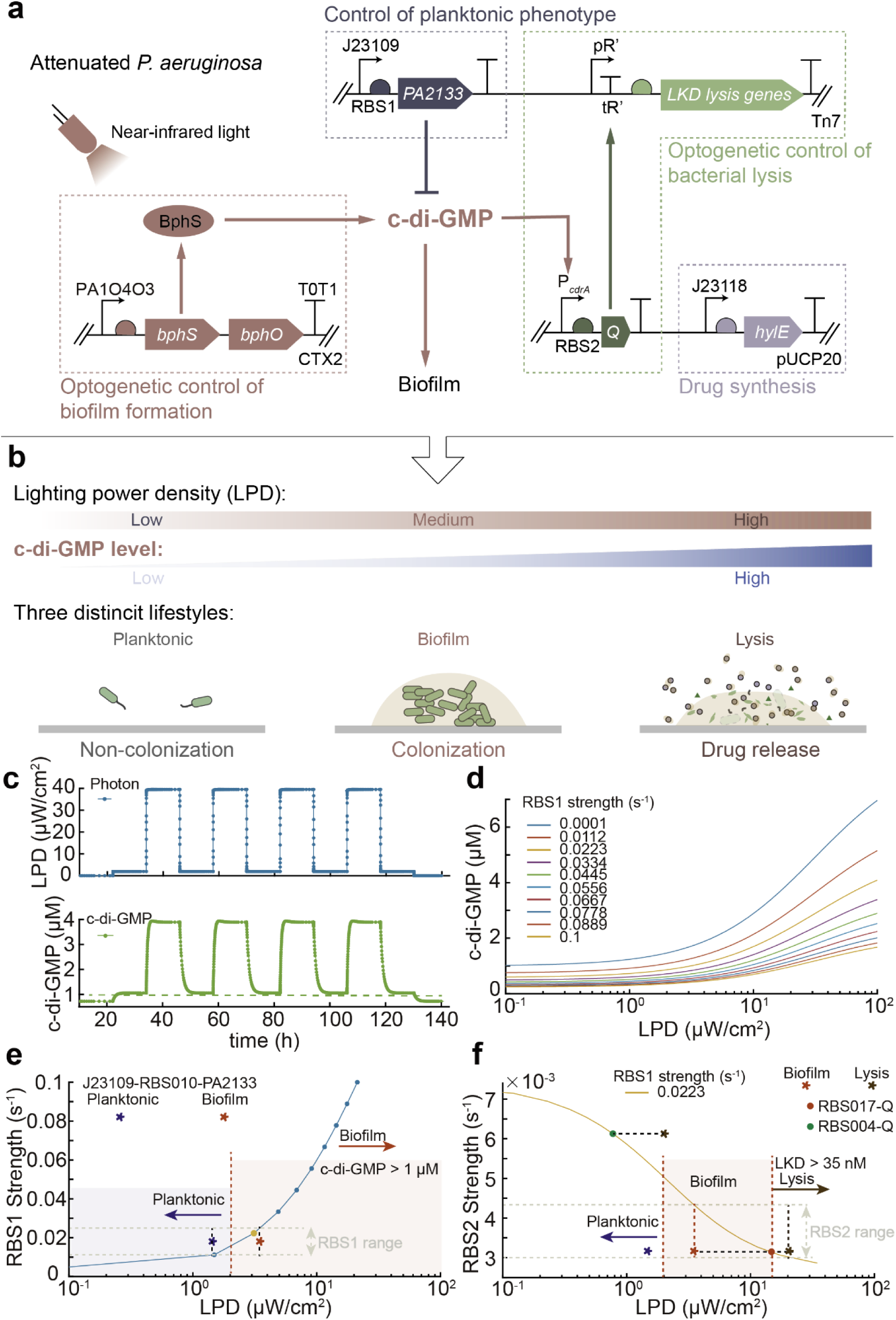
Genetic circuit design for programming bacterial lifestyles. (a) The programmable bacterial lifestyle system consists of four modules: the core NIR light responsive module producing c-di-GMP and promoting biofilm formation (red); a lysis circuit module driven by c-di-GMP-responsive promoter facilitating drug release (green); c-di-GMP hydrolysis module inhibiting bacteria from adhere to the surface (dark blue); and drug synthesis module to produce anticancer proteins (purple). (b) Schematic diagram of programmable manipulation of bacterial lifestyles using optogenetics. By adjusting the lighting power density (LPD) of near-infrared light (NIR), bacteria exhibit three different lifestyles: planktonic lifestyle (non-colonization), biofilm lifestyle (colonization) and lysis lifestyle (drug release). (c) The bacterial lifestyle transitions were programmed by manipulating intracellular c-di-GMP level through altering the LPD of NIR. (d) The level of c-di-GMP increases with LPD, and the profile correlated with the expression of *PA2133*. (e) The simulation results show the relationship between the threshold of LPD for biofilm formation and the intensity of RBS in front of *PA2133*. The threshold of c-di-GMP for biofilm formation was set at 1 μM in the model. When the intracellular c-di-GMP concentration was greater than 1 μM, the bacterial lifestyle changed from planktonic to biofilm. * indicates the experimental results to calibrate the kinetic constants in the model, and the color represents the bacterial lifestyle. (f) The RBS in front of *PA2133* was fixed when simulating the effect of *Q* gene expression level on the critical LPD threshold for bacterial lysis. The dots represent engineered strains with different RBS in front of *Q*.

### Modeling of dynamic programming bacterial lifestyle transitions

The genetic circuit for programming bacterial lifestyle transitions consists of four modules involving 22 different species and 30 chemical reactions (Table S4-6) following the mass action kinetics. In order to define an optimal strategy for subsequent high-throughput construction and screening of engineered bacteria, we developed a simulation model based on a chemical reaction network (CRN) to quantitatively characterize the genetic circuit for bacterial lifestyle programming (Fig. S4a). Firstly, we set the key parameters of the model according to previous studies and performed a parameter calibration in order to represent the real behaviors of each species in the experiment. We optimized the parameters in the photon activation and c-di-GMP signaling network (Table S7) to test whether the changes of model behavior were consistent with the experimental data, based on the analysis results of the effects of PA2133 expression at the threshold level of LPD for promoting bacterial biofilm formation (Fig. 1e). The modeling results indicate that the concentration of c-di-GMP in bacteria increases with LPD, which is consistent with data from the literature[53]. Moreover, we found that increased expression of *PA2133* by replacing promoter J23109 with J23105 inhibits bacterial colonization on surfaces, thus requiring a higher LPD level to promote biofilm formation (Table S8). The simulation results with the calibrated parameters (Table S7) show that bacterial lifestyle can be dynamically programmed by hierarchical modulation of the LPD of NIR (Fig. 1c). Furthermore, we used this model to investigate how these species change with different experimental parameters of the model (Fig. S5). By analyzing the effect of alterations of each parameter within an appropriate range on the model behavior, we found that the LPD range for maintaining bacterial biofilm lifestyle was jointly determined by the expression levels of *PA2133* and *Q* (Fig. 1f).

### Programming bacterial lifestyle transitions via hierarchically regulating the LPD of NIR

Our first goal was to obtain the strains displaying diverse lifestyles within a reasonable range of LPD through screening. In order to generate a series of strains with different expression levels of *PA2133* and *Q*, we constructed and characterized a ribosome binding sites (RBS) library with over a 400-fold dynamic range in *P. aeruginosa* (Table S10 and Fig. S6). Afterward, a self-made portable 96-well illuminator (Fig. S7) was employed to test the influence of LPD of NIR on the lifestyle of the engineered strains in a high throughput manner. Just as expected, PAO1 does not respond to NIR (Fig. S8). The strains capable of biofilm formation and lysis, which are distinctive features of our targeted strains, were identified using crystal violet staining and OD600 measurement (Fig. S10-S11). After multiple iterations, we successfully obtained the strain carrying the genetic circuit, wherein *PA2133* and *Q* gene are controlled by RBS010 and RBS017 respectively. This strain, denoted as H107, exhibited three distinct LPD-dependent lifestyles (Fig. 2a).

**Figure 2.**
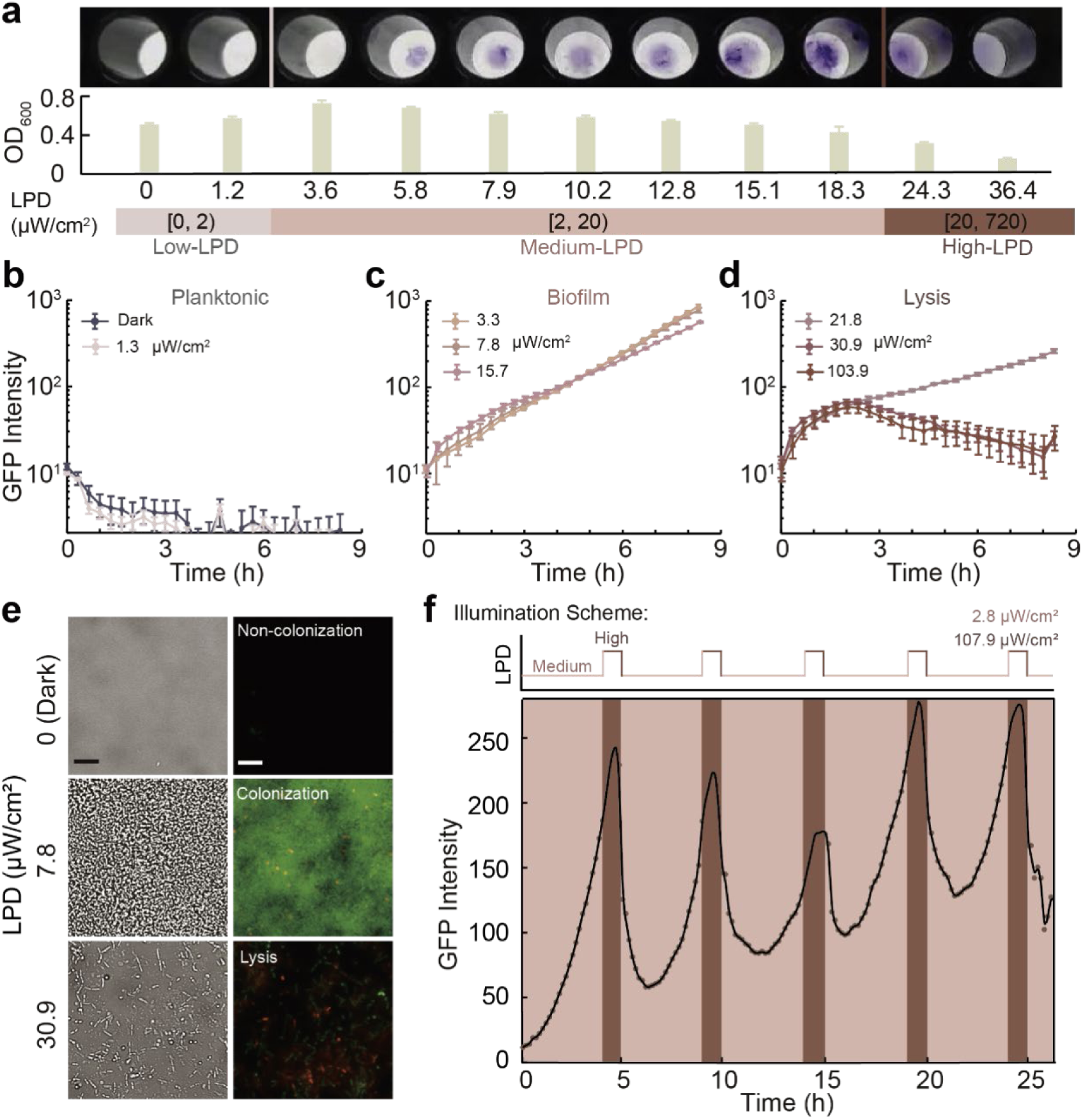
Programming bacterial lifestyles via hierarchically regulating the LPD of NIR. (a) Phenotypic characterization of engineered strains reveals the influence of LPD of NIR on bacterial lifestyle. Bacteria cultured statically in a 96-well plate were irradiated form the bottom with red light of different LPDs (0-720 μW/cm^2^) for 12 hours at room temperature. Crystal violet staining results(upper), OD_600_ of the culture supernatant (center) and corresponding LPDs applied (lower). LPD is divided into 3 intervals (denoted as Low, Medium and High-LPD), which were used to manipulate three bacterial lifestyles respectively (Planktonic, Biofilm and Lysis lifestyle). (b-d) Fluorescence profiles of H017 on the surface of microfluidic devices treated with Low-LPD (b), Medium-LPD (c) and High-LPD (d) of NIR. (e) Representative BF and fluorescence images of H017 grown for 8 h in the microfluidic device under comparative Low-, Medium- or High-LPD following PI staining. (f) Illumination scheme applied (upper) and the resultant fluorescence profile of H017 on the surface of microfluidic devices (lower). Fluorescence profile showed ‘biofilm-lysis’ oscillations along with repeated input signal of ‘Medium-High’ LPD switch. Symbols and lines in (b) to (d) and (f) represent the original and smoothing data, respectively. Scale bars for all images are 10 μm. Error bars represent standard error of mean (SEM) arising from ≥3 replicates.

To characterize the dynamics of lifestyle transitions of H017 irradiated with NIR of different LPDs, we monitored bacterial cell density on the surface of microfluidic devices using fluorescence microscope (Fig. S12). We observed that when illuminated with relatively Low-LPD (≤ 2 μW/cm^2^), it was difficult for H017 cells to colonize the surface (Fig. 2b and Movie S1). In contrast, under Medium-LPD (2 - 20 μW/cm^2^) H017 rapidly divided in situ to form micro-colonies and then developed into biofilms in 8 h (Fig. 2c and Movie S2). Under High-LPD (≥ 20 μW/cm^2^) lysis was observed (Fig. 2d, and Movie S3). Three lifestyles formed on microfluidics was further verified by 3D fluorescence imaging of bacterial cells after PI staining (Fig. 2e). The fluorescence intensity profiles of H017 treated with High-LPD suggested that the photon-activated bacterial lysis system has a response time of over 4 h and that the bacterial lysis behavior is asynchronous and lasts for over 10 h. To monitor the cellular levels of Q and c-di-GMP, P*cdrA*-gfp[55], a plasmid-based transcriptional reporter of c-di-GMP, was introduced into our engineered strains. The cellular levels of Q predicted by the computational modeling are consistent with our observations that the reporter fluorescence intensity increased with LPD and decreased after bacterial lysis. The wide distribution of fluorescence intensity within bacteria (Fig. S13) indicates that the expression noise of protein Q contributes to asynchronous lysis behavior and that bacteria containing high concentrations of Q lyse first[56].

To achieve the dynamic regulation of lifestyle transitions of H017 as predicted by the model, we optimized the illumination scheme in specific experiments. To begin with, 4-hour NIR with Medium-LPD (2.75 μW/cm^2^) is utilized to promote biofilm formation; then 1-hour NIR with High-LPD (107.91 μW/cm^2^) is used to induce bacterial lysis for 1 h. The entire lysis process takes about 6 hours. With a shorter period of NIR illumination with High-LPD, some bacteria will not undergo the lysis process. These surviving bacteria will regrow and form biofilm again (Fig. S14). This ‘biofilm-lysis’ lifestyle transition can be repeated cyclically under periodic NIR illumination of 4-hour Medium-LPD and 1-hour High-LPD (Fig. 2f and Movie S4). The results above suggest that the LPD of NIR employed to program the bacterial lifestyle transitions is tunable within the range of ‘μW/cm^2^ ’ level, which would greatly increase the availability of clinical treatments for deep-seated tumors.

### Programming bacterial lifestyle transitions facilitates the controllable release of therapeutic agents

To validate the role of the three bacterial lifestyles in tumor therapy in vitro, we co-cultured fixed human lung cancer cells A549 with our engineered bacteria in a microfluidic device (Fig. S15), and irradiated specific cancer cells with NIR of different LPDs. The illumination area can be determined according to the contour of the A549 cells with an established protocol (Fig. 3a). We observed that under NIR irradiation with Medium-LPD (7.72 μW/cm^2^) A549 cells were completely covered with biofilms that could resist the flow-induced shear (Fig. 3b), whereas in the absence of NIR illumination, bacteria cannot colonize the surface of A549 cells (Fig. 3c). These results suggest that the formation of biofilm facilitates the strong bacterial adhesion of the bacteria on the surface of the A549 cell. Thus, we speculated that biofilm formation in tumor sites may promote colonization. In addition, the planktonic lifestyle of therapeutic strains may contribute to the non-colonization of bacteria on normal tissues and decreased systemic cytotoxicity.

**Figure 3.**
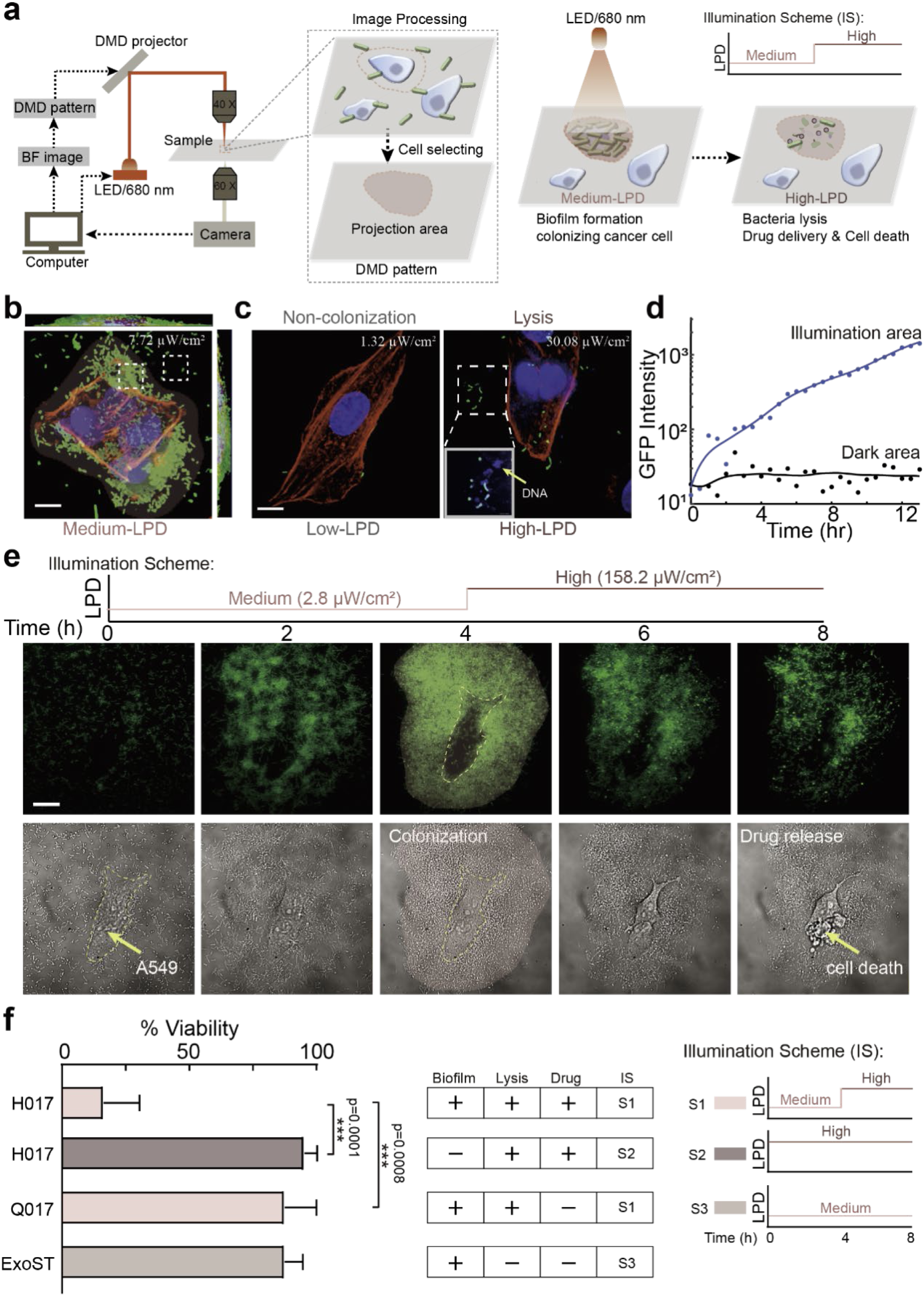
Programming ‘biofilm-lysis’ lifestyle transitions facilitates controllable release of therapeutic molecules. (a) Schematic of the experimental workflow of programming bacterial lifestyles on the surface of targeted human lung cancer cells A549 using microscope. The BF images of the cancer cells on microfluidics were acquired by a 60 × oil objective and then the profiles of selected cells were projected onto the cover glass by using a digital micromirror device-based (DMD) light- emitting diode (LED) projector through a 40 × air objective. The intensity of LED was controlled by MATLAB in real-time. Schematic diagram of programming ‘biofilm-lysis’ lifestyle transition in the targeted area by applying an illumination scheme. (b) Confocal projections of biofilms formed by RecA-H036 (labeled with GFP) co-cultured with fixed A549 cells under Medium-LPD of NIR (680 nm) for 8 h, and then stained with phalloidin (FITC, red) and DAPI (blue). (c) Laser scanning confocal micrographs of RecA-H036 co-cultured with fixed A549 cells under comparative Low- and High-LPD of NIR. Images are representative of NIR light area of the bacterial lifestyles monitored throughout the duration of the experiment, including three lifestyles: no attachment of cells to A549 cells, biofilms formation on A549 cells and bacterial lysis. The applied LPDs are shown in the upper right of the picture. Scale bar, 10 μm. (d) Fluorescence profiles of bacteria in illumination area or dark area as depicted in (b) (white dashed squares). (e) Schematic diagram of illumination scheme applied (upper) and the resultant time course of BF and fluorescence images of H017 co-cultured with A549 live cancer cells for 8 h in the microfluidic device, sequentially visualizing H017 biofilm formation, lysis and A549 cancer cell death. Scale bar, 20 μm. (f) Percentage viability of A549 live cells. A549 live cells co-cultured with H017 and treated with illumination scheme S1 to promote biofilm formation, bacterial lysis and drug release; co-cultured with H017 and treated with S2 to generate direct bacterial lysis and drug release; co-cultured with Q017 and treated with S1 to promote biofilm formation and bacterial lysis; co-cultured with ExoST and treated with S3 to promote biofilm formation. ***P<0.001, unpaired two-tailed t-test. Error bars represent SEM arising from ≥3 replicates.

To verify the benefits of bacterial lifestyle transitions in tumor therapy, we monitored the density of the bacteria in the illumination area with fluorescence microscope (Fig. 3d) and recorded the survival ratio of cancer cells co-cultured with engineered bacteria using different illumination schemes (Fig. 3e). We found that the density of the bacteria in the illumination area is proportional to the duration of NIR illumination with Medium-LPD (Fig. S15e). Thus, we hypothesized that programming the process of bacterial lifestyle (biofilm-lysis) transition could enable the controllable release of therapeutic agents into TME. To test this hypothesis, an illumination scheme (Fig. 3b) is adopted wherein NIR with Medium-LPD (2.75 μW/cm^2^) was applied for the first 4 hours, and then LPD was raised to 158.22 μW/cm^2^ for the next 6 hours. We observed that the survival ratio of A549 cells co-cultured with H017 was less than 20% (Fig. 3b and Movie S5), and there was no significant difference in the viability and morphology of A549 cells without NIR illumination. In this study, cell death can be determined by morphological changes and verified by PI staining (Fig. S16). Nevertheless, cell viability is slightly affected by direct lysis of H017 without biofilm formation (Fig. 3c and Movie S6), lysis of Q017 after biofilm formation without drug release (Fig. S17), and biofilm formation of ExoST without drug release (Movie S7). These results suggest that the localization, timing and dosage of therapeutic agents in the process of BMCT can be precisely manipulated according to our needs by programming bacterial lifestyle transition.

### Programming bacterial lifestyles transitions in solid tumors by NIR

To investigate whether the lifestyle of H017 in vivo can be programmed by modulating the LPD of NIR and whether this is consistent with our characterization in vitro (Fig. 4a), we irradiated A549 tumor-bearing mice according to various NIR illumination schemes after intratumoral injection of H017. We firstly re-calibrated the specific values for NIR with Medium-LPD and High-LPD respectively (Fig.4b), based on the tissue penetration efficiency of NIR[57]. The intratumoral bacteria is visualized using In Vivo Imaging System (IVIS). The fluorescence intensity of tumor tissues injected with H017 under dark conditions for 3 days (D3) was indistinguishable from that of PBS-injected controls (Fig. 4c, 4d, p=0.5737), suggesting that H017 could not colonize tumor tissues without NIR illumination. Tumors under NIR illumination with Medium-LPD (1.2 mW/cm^2^) for 3 days (M3) exhibited a higher bacterial fluorescence intensity than that of D3 (Fig. 4c, 4d, p=0.0028), indicating a higher density of bacterial colonization. Besides, the confocal fluorescence images of sections of tumors treated with M3 revealed that biofilms were formed in tumors (Fig. 4e, 4f).

**Figure 4.**
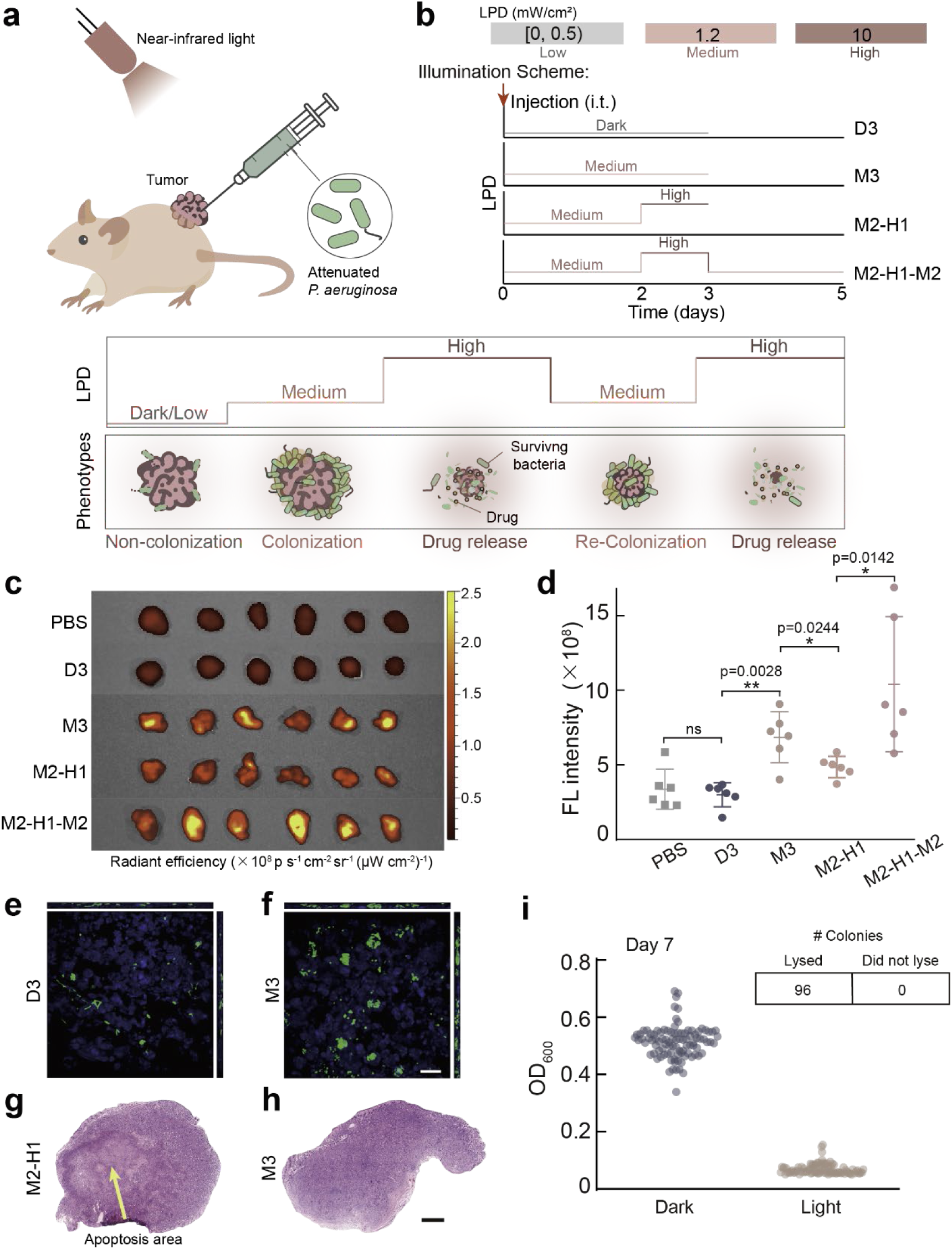
Programming bacterial lifestyles in solid tumors by NIR light. (a) Schematic diagram of dynamic programmable manipulation of intratumoral bacterial lifestyles for cancer therapy. Six to eight-week-old female BALB/c nude mice (n=6 per group) were subcutaneously (s.c.) inoculated with 10^8^ A549 cells into the flank. When tumor volumes reached about 100 mm^3^, mice were divided into five treatment groups. Group PBS received i.t. injection with PBS and cultured normally. The rest 4 groups received i.t. injection with 5×10^7^ CFU H017 and then treated with various illumination schemes as followed: group D3 was kept in dark conditions for 3 days, group M3 was illuminated with Medium-LPD for 3 days, group M2-H1 was illuminated with Medium-LPD for 2 days and high-LPD for 1 day successively, group M2-H1-M2 was treated with M2-H1 firstly and then illuminated with Medium-LPD for another 2 days. (b) Schematic diagram of experimental illumination schemes and specific values of each LPD. i.t. injection time point was depicted by red arrow. (c) IVIS images of tumors after treatment. (d) Fluorescence intensity of intratumoral bacteria determined by IVIS. Each dot in the plot diagram represents one counted tumor. Representative confocal microscopy image of frozen tumor sections (15 µm-thick) taken from mouse in group (e) D3 and (f) M3. DAPI (blue), H017 (green). H&E stained sections (8 µm-thick) of tumor tissues taken from mouse in group (g) M2-H1 and (h) group M3. (i) Distribution of OD_600_ values of bacterial supernatant originated from 96 single colonies cultured under darkness or a duplicate cultured with high-LPD illumination for 6 h. The colonies were harvested from tumors after treatment under Medium-LPD red light for 7 days. The grid above shows the number of successful lysis events. **P<0.01, *P<0.05, ns, not significant, unpaired two-tailed t-test. Error bars represent SEM arising from ≥3 replicates.

Next, we sought to assess whether the lifestyle transitions of intratumoral bacteria could be dynamically programmed by adjusting the NIR illumination scheme in vivo over a few days. We found that the fluorescence intensity of tumors treated with M2-H1 was lower than that of tumors treated with M3, indicating that bacterial lysis was triggered by 1-day High-LPD (10 mW/cm^2^) NIR illumination. Moreover, tumors treated with M2-H1 and stained with hematoxylin and eosin displayed cellular necrosis (Fig. 4g), whereas tumors treated with M3 maintained vitality (Fig. 4h). We speculate that lysis of bacteria within biofilm in the tumors treated with M2-H1 leads to sufficient HlyE release (Fig. S18). To verify whether the remaining bacteria after High-LPD illumination can grow and form biofilm again, we compared the fluorescence intensity of tumors treated with M2-H1-M2 and M2-H1 (Fig.4d, p=0.0142). The results showed that biofilm gradually reformed under another 2-day Medium-LPD of NIR illumination. Moreover, the intratumoral bacterial lifestyle transitions can be programmed with quite low LPD of NIR compared to that of used in PTT, making it promising for further clinical application for safety concerns. To confirm the long-term stability of our designed inducible lysis module after bacterial colonization in tumors, the growth of the bacterial colonies extracted from within tumors under 7-day NIR illumination with Medium-LPD (1.2 mW/cm^2^) was tracked under darkness and High-LPD of NIR illumination, respectively. We observed that colonies all lysed under NIR illumination with High-LPD (Fig. 4g), indicating that NIR sensitivity and lysis capacity were maintained in vivo.

### Programming the intratumoral bacterial lifestyle transitions enhances therapeutic efficacy

Given the ability to release antitumor drug HlyE in tumor grafts using NIR, we utilized H017 as a delivery vector to inhibit tumor growth (Fig. S19). Under 20-day NIR illumination with High-LPD, the tumor damage in the A549 tumor-bearing mice was determined by HE and TUNEL staining assays (Fig. S20). The mass of tumors with multiple injections of H017 under NIR illumination with High-LPD was significantly less than that of the control group (under darkness). In addition, the growth rate of tumor injected with ExoST under NIR illumination with High-LPD was indistinguishable from that observed under darkness, suggesting that the effect of biofilm formation and NIR illumination on tumor growth is negligible (Fig. S21). These results show that H017 was a highly efficient drug delivery system that can inhibit the growth of subcutaneous solid tumors in vivo. Though multiple injections of H017 followed by NIR illumination with High-LPD can inhibit tumor growth, excessive injections are not convenient and prone to side effects.

To investigate whether optimizing the process of bacterial lifestyles transitions enhances therapeutic efficacy, we monitored the relative tumor volume variation of the tumor treated with a single intratumoral injection of H017 and then illuminated the tumor using different illumination schemes as shown in Fig. 5a. Compared to other illumination schemes, tumor activity in mice injected with H017 using the illumination scheme (M2-H1) ×2 was significantly reduced over a period of 6 days (Fig. 5b-d and Fig. S22). Moreover, there was no significant difference in tumor volume between A549-bearing mice injected with Q017 using the illumination scheme (M2-H1) ×2 and those treated with ExoST using the illumination scheme M3×2 or D3×2 (Fig. 5d), indicating that neither natural cellular contents of our chassis bacteria nor biofilm formation of our chassis bacteria had a therapeutic effect on solid tumors.

**Figure 5.**
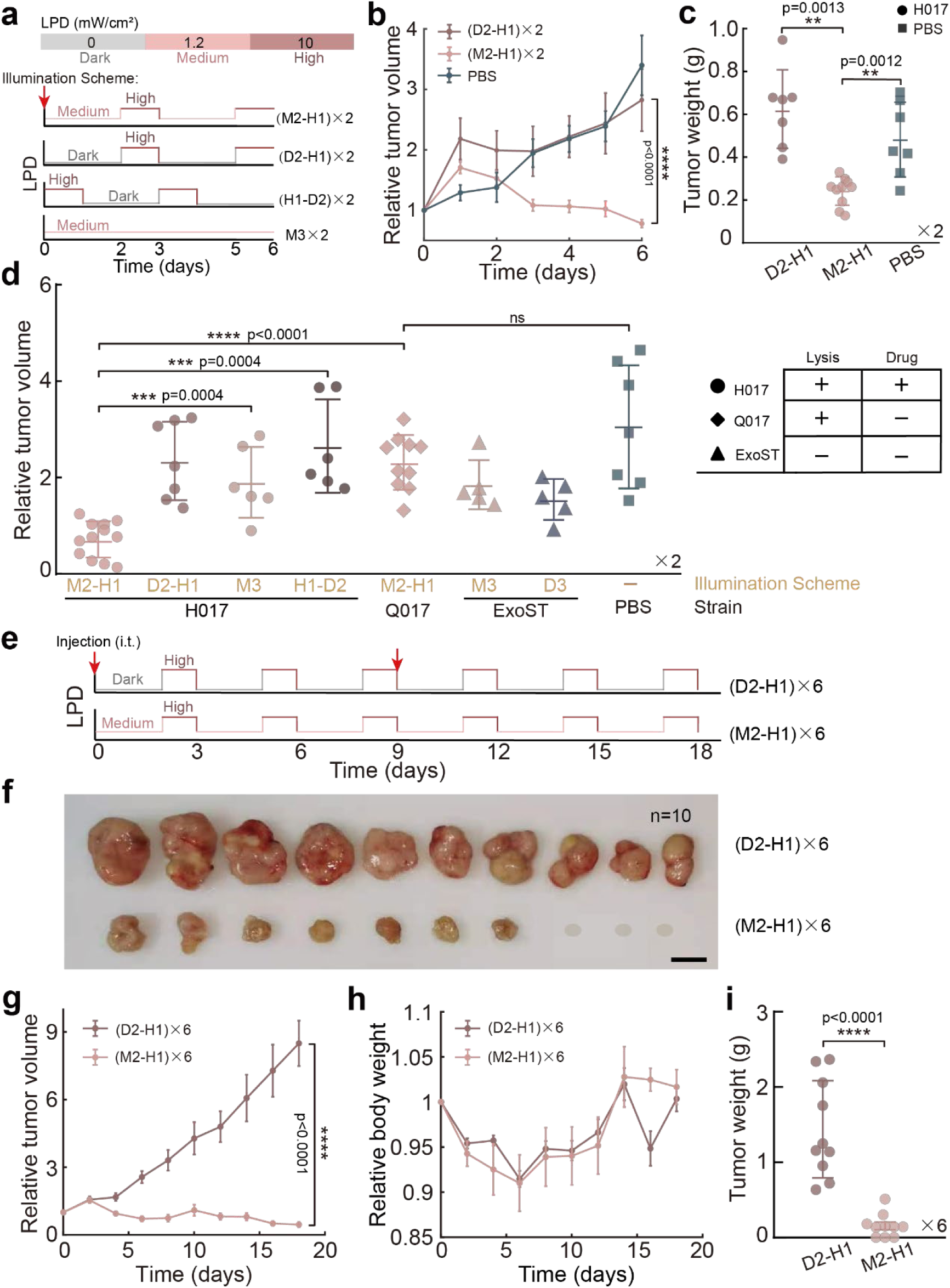
Tumor regression in mice induced by programming intratumoral bacterial lifestyles. Six to eight-week-old female BALB/c mice (n=5-12) were subcutaneously (s.c.) inoculated with 10^8^ A549-mCherry cells into the flank. When tumor volumes reached about 100 mm^3^, experiments were conducted. (a) Schematic diagram of experimental illumination scheme. All abbreviations have been mentioned except for D2-H1 and H1-D2. D2-H1 means culturing in dark conditions for 2 days and illuminating with high-LPD for 1 day, H1-D2 means illuminating with high-LPD for 1 day and then culturing in dark conditions for 2 days. ×2 means the process was repeated for 2 cycles. Time points of i.t. injection of 5×10^7^ CFU engineered bacteria (H017, Q017, ExoST) or PBS were depicted in red arrows. (b) Relative tumor volume over time and (c) distribution of weight of tumors. A549-mCherry tumor-bearing mice received i.t. injection of H017 were treated with illumination scheme (D2-H1) ×2 or (M2-H1) ×2, mice treated with i.t. injection of PBS and incubated normally were used as a control. (d) Distribution of relative tumor volumes. A549-tumor-bearing mice received i.t. injection of H017 and treated with (M2-H1) ×2, (D2-H1) ×2, M3 ×2, (H1-D2) ×2, or i.t. injection of Q017 and treated with (M2-H1) ×2, or i.t. injection of ExoST and treated with M3×2, D3 ×2, or i.t. injection of PBS and incubated normally. Functional compositions of each engineered strains were also depicted in right picture. (e) Schematic diagram of illumination schemes applied, ×6 means the process was repeated for 6 cycles. Time points of A549-mCherry tumor-bearing mice received i.t. injection of 5×10^7^ CFU H017 were depicted by red arrows and (f) images of the resultant tumors (scale bar, 1 cm). The gray ellipse indicates that the tumor has been completely eliminated. (g) Relative tumor volume and (h) relative body weight of A549-mCherry tumor-bearing mice over time. (i) Distribution of weight of tumors. ****P<0.0001, ***P<0.001, **P<0.01, ns, not significant, unpaired two-tailed *t*-test. Each dot in the plot diagram represents one counted tumor. Error bars represent SEM arising from ≥3 replicates.

Considering that our designed ‘biofilm-lysis’ transition can promote bacteria recolonization after lysis, sustained and controlled drug release is possible if an appropriate illumination scheme is adopted. To improve the therapeutic efficacy, we extended the illumination scheme to 3 cycles and repeated the intratumorally injection-illumination treatment (Fig. 5e, 5f, 5i). We found that 30% of the tumors disappeared after treatment and the growth of the remaining tumors was significantly inhibited with double injections and multiple cycles of NIR illumination. It should be noted that no significant adverse effects were observed (Fig. 5h). Overall, our in vivo experiments indicated that complete tumor eradication can be achieved by programming bacterial lifestyle transitions with an optimized illumination scheme.

## DISCUSSION

We have developed a programmable bacterial lifestyle transitions system based on NIR light for the controllable release of antitumor drugs into TME and exemplify a methodology for promoting bacterial colonization of tumor sites. Clinical studies have shown that increasing tumor-specific colonization by bacteria can greatly enhance therapeutic efficacy and reduce toxicity to normal tissues. In this study, the formation of biofilm in tumor tissue significantly enhanced bacterial colonization. High levels of bacteria were observed to persist within tumors for at least 7 days (Fig. 4e), thereby releasing more therapeutic agents for better therapeutic efficacy (Fig. 5c). The formation of *P. aeruginosa* biofilms mediates long-time lung colonization. And the synthesis of extracellular polymeric substances (EPS) forms sticky and tangled fibers that connect bacteria cells to each other and to various surface[58]. Thus, we speculate that the biofilm lifestyle acts as a defense barrier that protects the bacteria embedded in the EPS matrix against innate host defense[59] and shear caused by fluid flow[60]. In addition, EPS produced on the surface of cancer cells inhibits the adhesion of cancer cells to endothelial cells, resulting in metastasis disruption[42, 61]. Also, c-di-GMP which regulates biofilm formation, motility and virulence, is a promising candidate as antitumor drug by stimulating inflammation via STING[62]. While the above hypotheses require further validation, our results indicated that biofilms have shown great application potential in BMCT.

Besides, optogenetics provides a promising approach for gene- and cell-based therapies via precisely manipulating various cellular activities with high spatiotemporal resolution[63]. And the deep tissue penetration of NIR light offers a non-invasive treatment modality for a variety of diseases[64, 65]. The illumination intensity and pattern of multiple light spots, especially lasers, can be easily controlled by computer algorithms, enabling neuroscientists to stimulate hundreds of precisely targeted neurons simultaneously[66]. In this study, we programmed intratumoral bacterial lifestyle transitions by modulating the LPD of NIR, opening up the possibility of customized cancer therapy. We can use the specialized device to steer near-infrared lasers to precisely target bacteria within deeper tumors and design a personalized treatment regimen by changing the illumination scheme according to the size of the tumors and the type of antitumor drugs.

In addition to programming the intratumoral bacterial lifestyle, we also engineered the extratumoral bacterial lifestyle, which is uncommon and difficult to achieve in the current BMCT. The therapeutic strains in our system, which continuously express PA2133, produce a relatively low level of c-di-GMP intracellularly and cannot colonize various tissue surfaces under darkness (Fig. 3a). This is also confirmed by the result that the fluorescence intensity of tumors injected with H017 under darkness for 3 days was essentially indistinguishable from that of tumors injected with PBS (Fig. 4c). The induction of PA2133 (c-di-GMP hydrolysis module) results in the maintenance of planktonic lifestyle which exhibits high susceptibility to antimicrobials[67], allowing *P. aeruginosa* biofilms to disperse and thus be eliminated by the host immune system[68]. Furthermore, the engineered bacteria can initially adhere to surfaces within 2 h (Fig. 2b), which is necessary for bacterial colonization within tumors. In summary, programming lifestyle transitions of the engineered bacteria according to various environments would greatly improve safety while enhancing the antitumor efficacy Our results indicate that bacterial lifestyle can be easily switched between two or more lifestyles by modulating the NIR illumination scheme in vitro. With our designed genetic circuit, bacteria can achieve multiple cycles of ‘biofilm-lysis’ transition, resulting in sustained release of drugs to eliminate tumors. However, the therapeutic efficacy gradually diminished after three cycles of the illumination scheme M2-H1 in vivo. Some tumors disappear completely with a second injection of the engineered bacteria (Fig. 5f). The possible reasons for the above results include: 1) The loss of the plasmid in the bacteria containing the lysis trigger gene *Q* and the mutation of bacteria leads to the failure of the lysis system; 2) Due to the limitations of experimental conditions, we were unable to obtain the best illumination scheme by observing the bacterial density within the tumor in situ. Further improvements would derive from strategies for longer-term circuit stability and the utilization of additional therapeutic agents.

## MATERIALS AND METHODS

### Strains and Mammalian cells culturing

The chassis strain used in this study was PAO1, and bacterial strains used in this study are listed in Table S11. Generally, strains were grown in either LB broth or minimal medium for *Pseudomonas* (FAB) at 37 ℃ and supplemented with antibiotics as and when required. Before each assay, cells cultures were inoculated from a single colony and grown in the dark at 37 ℃ until the mid-exponential phase (OD600 ≈ 0.8), unless otherwise specified. Mammalian cells A549-mCherry and A549 were obtained from the Cell Resource Center, Peking Union Medical College (which is the headquarter of National Infrastructure of Cell Line Resource, NSTI). Cell lines were cultured in Mccoy’s 5A medium (KeyGENE BioTECH) supplemented with 10% fetal bovine serum (FBS, Gibco) and penicillin/streptomycin (Sangon Biotech), placed inside the incubator at 37 ℃ with 5% CO2. When A549 cells covered 80%-90% surface area of the dishes, they were washed with PBS and dis-adhered with Trypsin-EDTA solution (Sangon Biotech). Cells were then collected by centrifugation and resuspended in McCoy’s 5A medium + FBS or PBS for further use.

### Construction of plasmids and PAO1 derivatives

To generate an attenuated strain of *P. aeruginosa*, the following virulence factors genes were sequentially deleted from the chromosome of the wild-type strain PAO1: *vfR*, *exoS*, *exoT*. The core NIR light responsive module *bphS* was integrated into the chromosomal *attB* site by using plasmids miniCTX2. The c-di-GMP hydrolysis module *PA2133* and lysis cassette *LKD* was cloned into vector miniTn7, and then inserted into the genome *attTn7* site. Anticancer drug synthesis gene *hlyE* and antiterminator *Q* for activation of the lysis system in response to c-di-GMP concentration was cloned into vector pUCP20We also constructed and characterized the RBS library in PAO1 cells. All plasmids were constructed using basic molecular cloning techniques and Gibson assembly. Additional details for construction of the optogenetic and reporter strains are given in Supplementary text.

### Computational Model

Using a chemical reaction network (CRN) representation, we describe and quantify the dynamics of the designed genetic circuit composed by 22 different species (Table S6) and 30 chemical reactions (R1 to R30 in Table S4) that follow the mass action kinetics. The diagram of model was shown in Fig. S4 and the simulation was computationally implemented using a MATLAB toolbox (Simbiology). The parameters of reactions in this model, obtained from previous literature reports or our experimental results, were list in Table S7. The detailed computational model derivation is described in the Supplementary text.

### High-throughput screening assay

Based on RBS library, a series of light-responsive strains can be constructed by synergistically altering the gene expression of PA 2133 and Q. The engineered strains were then incubated in 96-well plate supplied with different the lighting power density (LPD) by our designed device (Fig. S10). The threshold of LPD for cell lysis was determined by measuring the OD600 of the supernatant with a microplate reader. *P. aeruginosa* biofilm was formed on 96 well black bottom plates (Corning) using a modified microtiter plate crystal violet assay that is a useful method for the growth and analysis of biofilms in static systems.

### Microfluidic Experiments and Microscopy

Our microfluidic devices (Fig. S15a) were made of polydimethylsiloxane (PDMS) using a well-established protocol. The device is composed of one bacteria/cell port, one waste port, one main channel and several chambers (300 μm × 300 μm × 100 μm) placed in both sides of the main channel. These chambers were used for bacterial culture or co-culture of bacteria and mammalian cells in microscopy experiments. All microfluidic experiments were conducted at 30 ℃ under darkness, and the flow rate was 0.5 mL/h. Bacterial strain to be verified was harvested at OD600 ≈ 0.6 cultured with FAB medium at 37 ℃, and diluted bacterial culture (OD600 ≈ 0.1) was injected into the microfluidic device. The device was then placed on XY-stage (Ludl) of the microscope and montage fields for LED (680 nm) illumination were selected manually through MATLAB. These fields were illuminated with different LPDs of laser under control of illumination scheme generated by MATLAB. Commonly, 8-10 fields were selected at a time, and the corresponding LPD can be changed from 0 to 158 μW/cm^2^. The invert fluorescence microscope (Olympus, IX71) equipped with a 60×oil objective and two sCMOS cameras (Zyla 4.2, Andor, 2048 × 2048 pixels) was acquire bright field (BF) images and fluorescence images (3 frames per hour). The surface attached bacteria were allowed to continuously grown in the microfluidic chamber and illuminated under 680 nm NIR light for 8 h.

Afterward, propidium iodide (PI, Sangon Biotech) dissolved in FAB medium (10 μg/mL) was injected into the channel to stain dead cells. Live cells (green fluorescence) and dead cells (red fluorescence) were excited using 488 nm and 561 nm lasers, respectively, and imaged in two emission channels (524 nm and 605 nm). Fluorescence intensity profiles were obtained by analyzing frames from the fluorescent channel and plotting the mean pixel intensity over time through a general image processing algorithm coded by MATLAB.

### Mammalian cells co-cultured with bacteria

All Bacteria and cancer cell co-culture experiments were performed in microfludic device supplemented with Mccoy’5A medium containing 10% FBS and 30 μg/mL gentamycin. The corresponding LPD for manipulating bacterial lifestyles vary slightly under different culture conditions. For culturing live cell in microfluidic devices, about 10^4^ resuspended A549 cells were injected into the microfluidic device with 1 mL syringe and cultured in the incubator at 37 ℃ with 5% CO2 overnight. Medium was replaced by fresh medium once after culturing for 6 h to remove dead cells and wastes. Then, the device was incubated overnight with the flow of 0.5 mL/hr. For preparation of fixed cells in microfluidic devices, adequate 4% paraformaldehyde was injected into the microfluidic devices loaded with live A549 cells to replace the cell culture medium. The A549 cells were fixed overnight at 4 ℃. On the next day, the device was placed on XY-stage of microscope and appropriate microscope fields were selected. After that, 1 mL of the diluted bacterial culture (OD600 ≈ 0.1) was injected into the microfluidic device loaded with live/fixed A549 cells and placed for 20 min without flow. Then, a flow with a constant flow rate (0.5 mL/h) provided by a syringe pump and a gas-tight syringe were applied to remove the unattached bacteria. A549 cell survival ratio can be determined by the change of cell morphology and verified by the results of PI staining. Bright field and fluorescence images are acquired under different illumination scheme.

### Characterization of the lifestyle of bacteria on A549 cells

The microfluidic device loaded with fixed A549 cells was injected with RecA-H036 strains. Then, BF images were acquired and converted from TIFF to RGB format. In one field, the profile of the fixed A549 cells was outlined through software ImageJ and a closed area was generated. The specific A549 cells can be irradiated with NIR light through a DMD projector (Mosaic Andor), and the illumination schemes was controlled by MATLAB. Phalloidin (Cytoskeleton) and DAPI Stain Solution (Sangon Biotech) was directly added into the channel of the microfluidic device for staining according to standard protocol if necessary. Multi-field and z-scanning fluorescence images (step size = 0.5 μm) of a tumor cell coated with RecA-H036 (Fig 3b) was acquired by a spinning-disc confocal microscope (IX81, Olympus). ImageJ was used to reconstruct the three-dimensional structure.

### Mouse Experiments

This study was approved by the Local Ethics Committee for Animal Care and Use (Permit Number: USTCACUC 1901026), and the experiments involving animals were used according to the animal care regulations of the University of Science and Technology of China. Sacrificed animals were euthanized by Cervical dislocation when the tumor size reached 2 cm in diameter or after recommendation by the veterinary staff. Animal experiments were performed on BALB/c nude mice (Charles River) that were allowed to acclimatize to the institutional animal facility for one week prior to experiments.

#### In vivo cytotoxicity assay

Bacteria cultured in FAB medium supplemented with 30 mM glutamate and 1 μM FeCl3 were spun down until exponential phase and washed 3 times with sterile PBS before injection into mice. Subcutaneous injections of bacteria were performed at a concentration of 5×10^8^ CFU bacteria per mL in PBS with a total volume of 100 μL injected per flank. After subcutaneous administration of PAO1- BphS-LKD-GFP or ExoST, mice were closely monitored and weighed daily for 8 days. The mortality and weight change of mice was calculated compared with day 0 (first day of treatment administration).

#### Subcutaneous tumor model

Six to eight-week-old female mice were subcutaneously (s.c.) inoculated with A549-mCherry or A549 cells into the flank. All pre-conditioned tumor cells were adjusted to 10^8^ cells per ml in PBS, and implanted subcutaneously at a volume of 100 μL per flank. Tumors were measured with a caliper every 2 days. Tumor volume (in cubic millimeters) was calculated using the following formula: 0.5×(L×W^2^), where L is the length, W is the width of the tumor in millimeters. Tumors were typically grown to an appropriate size (about 100 mm^3^) before experiments.

#### Biodistribution of bacteria in tumor

GFP-labeled *P. aeruginosa* H017 (5 × 10^7^ CFU per mice) were intratumorally injected into A549 tumor-bearing BALB/C mice. Then, the mice were randomly divided into five groups with six mice in each group (Fig. 4a). In the mouse experiment, the LPD of red light in the cage is adjusted by changing the number of LED lamp beads that emit red light around the cage, and the detailed method for controlling the illumination scheme are presented in the Supplementary text. Experiment groups illuminated by NIR with the various illumination schemes was injected with H017, and the control group was injected with PBS. At six days after the two cycles of illumination, the mice in each group were killed, and the tumor tissues were fixed with 4% (wt/vol) paraformaldehyde overnight at 4 °C and imaged by an IVIS in vivo animal imaging system to quantify the fluorescence intensity after mice treated with the NIR light. The tumors tissues were then dehydrated in 30% sucrose at 4 °C, and embedded in OCT before sectioning and staining. Frozen tissues used for immunofluorescence and Hematoxylin and eosin (H&E) staining were sectioned (8 μm), while the sections used for laser confocal fluorescent microscopy are 15 μm. Sections were stained with DAPI and then imaged using a spinning-disc confocal microscope equipped with a 100×oil objective and an EMCCD camera (Andor). The z-scanning (step size = 0.5 μm) confocal images were acquired to reconstruct the three-dimensional structure of bacteria biofilm (green) formed in tumor tissues with NIR illumination.

#### Antitumor efficacy

In order to test the effect of biofilm formation and bacterial lysis on tumor growth, the subcutaneous A549-mCherry tumor-bearing BALB/C mice were treated with various illumination schemes indicated in Fig.5a after injected with ExoST and Q017 strains respectively. The tumor volumes and body weight of mice were monitored every day. Relative tumor volume over time for mice was recorded to follow physical tumor growth. The tumor tissues were collected for image, weight and histology analysis.

### Statistical analysis

All qualitative images presented are representative of at least three independent duplicate experiments. Mice were randomized in different groups before experiments. All experiments data are expressed as the mean ± SEM. Statistical significance between control and treated groups was evaluated using two-tailed Student’s t-test using the statistical program of GraphPad Prism software. Differences between experimental groups were considered significant when P values were < 0.05. Statistical details of the experiments were included in the figure legends.

## ACKNOWLEDGEMENTS

We thank J.J. Wei for proofreading the manuscript.

## AUTHOR CONTRIBUTIONS

F.J. and RR.Z. designed the experiments. RR.Z., SW.F. performed experiments and analyzed data. YM.G. helped to construct bacterial strains. JR.X. constructed and characterized the RBS library. Y.L. designed and fabricated the 96-well illuminator. AG. X. helped to obtain microscope images. RR.Z., SW.F. L.P. and F.J. contributed jointly to data interpretation and manuscript preparation. All authors reviewed the manuscript.

