## Supplementary information for "Programming the lifestyles of engineered bacteria for cancer therapy"

#### Contents


| | |
|---|---|
| **Supplementary Information Text** | **2--16** |
| **Figures S1-S22** | **17--34** |
| **Tables S1-S11** | **35--42** |
| **Movies S1-S7** | **43--49** |
| **Supplementary References** | **50** |

### Supplementary Information Text:

**Supplementary Note 1: Evaluate the attenuation effects of *P. aeruginosa*.**

**1）In vitro cytotoxicity assay.**

For in vitro cytotoxicity assay, resuspended A549 cells were seeded in 96-well plates (Corning) at a density of about $10^4$ cells per well and grown at 37 ℃ with 5% $CO_2$ for 24 h before experiment. Bacterial strains were grown in FAB medium[1] supplemented with 30 mM glutamate and 1 μM $FeCl_3$ until exponential phase, washed twice with PBS, resuspended and diluted in Mccoy's 5A medium. The cytotoxicity assay in vitro was performed at a multiplicity of infection (MOI) of 50 in McCoy's 5A supplemented with 10% FBS. After 24 h infection, medium was removed from the wells, placed into microfuge tubes and spun for 2 min at 14000 rpm to pellet the bacteria and A549 cell debris. The level of lactate dehydrogenase (LDH) in the supernatant was then assayed in triplicate using a LDH Cytotoxicity Assay Kit (Beyotime). Cells treated with lysis buffer were used as a positive control for maximum LDH release, untreated cells were used to assess background LDH release.

**2）Annexin V-FITC/PI-staining.**

A549 cells were injected into flow cell[2] and cultured in Mccoy'5A media supplemented with 10% FBS, placed inside a tissue culture incubator at 37 ℃ maintained at 5% $CO_2$. Furthermore, 1 mL of the diluted bacterial culture ($OD_{600} \approx 0.1$) was injected into the flow cell channel and the device was placed for 15 min without flow. Then, a flow with a constant flow rate (3 mL/h) provided by a syringe pump (Harvard Apparatus, Holliston, Phd2000) and a gas-tight syringe (SGE, 25 mL) were applied to remove the unattached bacterial cells. A549 cells were stained using the FITC Annexin V/PI kit (Beyotime) which is a commonly used approach for studying apoptotic cells. Meanwhile, an inverted fluorescence microscope (IX81, Olympus) equipped with a 60× silicone oil objective and a sCMOS camera (Andor, Zyla 4.2) was used to acquire the BF and fluorescence images. The staining results show that the rounded cells are all cells that have begun to be apoptotic (Fig. S1 b). Therefore, we can calculate the survival ratio of cells based on the morphology of the cells in the following experiment.

**Supplementary Note 2: Optimization of the inducible lysis genetic circuit.**

**1) Construction of plasmids used for screening of effective lysis genes in PAO1.**

All promising lysis genes were list in Table S3. *AlpA*[3], *alpBCDE*[3], *PA0614*[4], *PA0629*[4] and *PA1150* were amplified from PAO1 genome, λ lysis cassette[5] was amplified from phage λ genome, the lysis cassette from *Pseudomonas* phage LKD16[6] named *LKD* for short was synthesized by Sangon Biotech, all PCR fragments were cloned into pJN105 and placed downstream of araC promoter.

**2) Characterization of lysis efficiency.**

The PAO1 single colonies (isolates) were inoculated into 3 mL liquid LB medium and shaken overnight at 37 °C, 250 rpm. Cells were harvested and washed twice with 300 mM sucrose, and then resuspended in 500 μL sucrose solution for further use. The constructed plasmids (100 ng) for screening lysis genes were introduced into PAO1 (100 μL per tubes) by electroporation. Then, 900 μL of fresh LB medium was supplied immediately and continue to culture for 1 h at 37 ℃. After that, 100 μL of bacterial culture of each tube was taken and spread over the surface of LB agar plate containing 30 μg/mL gentamicin. After overnight incubation, the number of isolates was counted. We found that bacteria electroporated with the plasmid *LKD*-pJN105 formed no isolates. Subsequently, we adjusted the RBS in front of *LKD* and found that only the plasmid without RBS in front of *LKD*, which is *LKD*-Remove RBS-pJN105, could form colonies on the LB agar plate after electroporation into PAO1.

**3) Improve the stability of inducible lysis module.**

Considering that *LKD*-pJN105 lyse PAO1 even at leaky expression level (Fig. S2 b), it was selected for further optimization. Though *LKD*-Remove RBS-pJN105 could induce lysis of PAO1 even without RBS upstream of the lysis cassette, it was unstable since the lysis system did not work after fourth generation of induction (Figure. S2 d). To minimize the leaky expression, we employed λ late gene expression system and *mScarlet* was used as an indicator. All these parts were cloned into vector miniTn7 for single copy genome expression and *Q*-pJN105 was used for induction of gene expression (Fig. S3). Our attempt turned out to be effective with merely no fluorescence detected in pR'-tR'-mScarlet. Though the fold change seems not high, it's enough for LKD to lyse PAO1. Nonetheless, the expression level of *Q* gene ought to be further optimized for lifestyle control. RBS upstream of *Q* and ssrA degradation tag of *Q* were screened altogether. The ASV peptide tag was finally selected since the others are too strong to trigger Q-mediate lysis.

**Supplementary Note 3: Construction and characterization of RBS library.**

Plasmids used for calibrating RBSs strength contain two modules: *sfGFP* driven by constitutive promoter was used as an internal control, and *cyOFP* driven by J23102 and RBS(N) (RBS to be test) was the reporter. Template plasmid J23105-B0034-*sfGFP*-T0T1-J23102-B0034-*cyOFP*-pUCP20 was firstly generated by PCR and Gibson assembly. To change the RBS upstream of *sfGFP*, a set of random primers were synthesized, all these primers have one 20 bp homology region for PCR amplification and another 25 bp homology regions for Gibson assembly, the 12bp RBS region was set to be NNNNNNNNNNNN (N=A, T, C, G). The resultant plasmids J23105-RBS(N)-sfGFP-T0T1-J23102-cyOFP-pUCP20 were finally obtained by Gibson assembly and electroporated into PAO1 to generate the RBS library.

To measure the RBSs strength, all bacterial strains in RBS library were cultured overnight and diluted 100× to fresh FAB medium supplemented with 30 mM glutamate, 1 μM $FeCl_3$ and 30 μg/mL gentamicin until $OD_{600} \approx 0.6$. The resultant cells were resuspended to $OD_{600} \approx 0.02$ and then loaded 10 μL bacterial cultures on agarose pads (FAB medium containing 30 mM glutamate, 1 μM $FeCl_3$, 30 μg/mL gentamicin and 1% agarose). When the liquid was about to dry, the pads were flipped and transferred to cover slips. Fluorescence images were acquired by a laser scanning confocal microscope. The RBSs strength was measured by the ratio of fluorescence intensities of *mScarlet* and *cyOFP* with laser scanning confocal microscope (Fig. S6 and Table S10).

#### Supplementary Note 4: Construction plasmids and PAO1 derivatives

**1）Construction of plasmids used for genome insertion.**

The λ late gene expression system pR'-tR' was amplified from λ genome[7] by PCR. *LKD* was placed downstream of pR'-tR' and RBS BBa_B0034 for minimizing leaky gene expression by overlap extension PCR. Phosphodiesterase encoding gene *PA2133* was obtained by PCR from PAO1 genome and under control of promoter J23109 and RBS RBS010 (from RBS library, see Table S10). Plasmid pR'-tR'-B0034-*LKD*-T0T1-J23109-RBS010-*PA2133*-Tn7 was finally cloned by Gibson assembly of these two PCR fragments and linearized vector pTn7-L(T0T1)[8]. To tune the expression level of *PA2133*, the upstream promoter or RBS of *PA2133* was replaced via Gibson assembly. The ligation product was transformed into commonly used chemically competent *E. coli* host Top10.

Plasmid PA1/O4/O3-*bphS*-T0T1-CTX2[9] was used as a template vector for insertion of the PCR fragment J23102-B0034-*sfGFP* to generate plasmid PA1/O4/O3-*bphS*-J23102-B0034-*sfGFP*-CTX2 via Gibson assembly. Plasmids pR'-tR'-B0034-*mScarlet*-Tn7 and pR'-B0034-*mScarlet*-Tn7 were cloned by Gibson assembly as well. The map of plasmids used for genome insertion are shown in Fig. S9.

**2）Construction of plasmids used for expression of Q and hlyE.**

Antiterminator protein encoding gene *Q* and hemolysin encoding gene *hlyE* were amplified from λ phage genome and *E. coli* Top10 genome by PCR respectively. P*cdrA*-*Q*-pUCP20 was cloned by replacing *gfp*(mut3) in P*cdrA*-*gfp*(mut3)-pUCP20 with *Q* via Gibson assembly. PCR fragment J23118-RBSII-*hlyE* together with terminator T0 and T1 was inserted into backbone P*cdrA*-*Q*-pUCP20 to generate plasmid J23118-RBSII-*hlyE*-T0T1-P*cdrA*-*Q*-pUCP20. To tune the expression level of *Q*, RBS upstream of Q and ssrA degradation peptide tag in P*cdrA*-*Q*-pUCP20 and P*cdrA*-*Q*-T0T1-J23118-RBSII-*hlyE*-pUCP20 were replaced by Gibson assembly.

**3）Construction of plasmids used for gene knocking out.**

The upstream and downstream flanking regions (~1000-1500 bp) of *vfr*, e*xoS* and *exosT* were obtained through PCR from PAO1 genome. The PCR fragments of each gene were inserted into the linearized pEX18Gm vector via Gibson assembly to generate plasmids pEX18Gm-*vfr,* pEX18Gm-*exoS,* pEX18Gm-*exoT.*

**4）Construction of PAO1 derivatives.**

All Tn7- and CTX2-based plasmids were sequenced and then inserted into PAO1 genome with standard protocols[10, 11]. The resultant strains include PAO1-BphS, PAO1-LKD, PAO1-BphS-LKD, PAO1-BphS-LKD-GFP, pR'-mScarlet, pR'-tR'-mScarlet, etc. The gentamicin and the tetracycline resistance flanked by FRT sites were removed by expressing Flp recombinase from plasmid pFlp2[12] .

ExoST was obtained by triple use of two-step allelic exchange[13] in PAO1-BphS-LKD-sfGFP with plasmids pEX18Gm-*vfr,* pEX18Gm-*exoS,* pEX18Gm-*exoT.* Taking the gene knockout of *vfr* as an example, the detailed experimental protocol is as follows. The recombinant plasmid pEX18Gm-*vfr* was electroporated into PAO1-BphS-LKD-sfGFP and the resultant strain was screened on LB agar supplemented with gentamycin and NaCl-free LB agar containing 15% sucrose successively. Mutations were finally verified through PCR and sequencing. The other two genes *exoS* and *exoT* were knocked out similarly. RecA was obtained by double knocking out of *vfr* and *recA* with the same protocol.

H017、H004、Q017、Q018 and all other similar ExoST derivatives for were obtained by electroporation of corresponding plasmids into ExoST. For example, ExoST containing plasmid P*cdrA*-RBS017-*Q*(ASV)-T0T1-J23118-RBSII-*hlyE*-pUCP20 was named H017 (H for *hlyE*, 017 for RBS017) and Q017 means ExoST containing plasmid P*cdrA*-RBS017-*Q*(ASV)-pUCP20 (017 for RBS017, without *hlyE*).

When required, antibiotics were added to medium at the following concentrations (μg/mL): gentamicin, 15, ampicillin, 100, tetracycline, 10 (*E. coli*); ampicillin, 300, gentamicin, 30, tetracycline, 100 (*P. aeruginosa*).

#### Supplementary Note 5: Modeling of the genetic circuit based on Chemical Reaction Network.

##### 1) System biology model of the programmable bacterial lifestyle system.

We used a MATLAB package (Simbiology) for simulation of biological networks. The diagram of computational model is shown in Fig. S4 a, and each step of the biological process can be represented by the corresponding chemical reaction (R1-R30). These reactions can be divided into 6 different categories (Table S4), including 1) photon activation of BphS (**R1** to **R3**), 2) c-di-GMP signaling (**R4** to **R10**), 3) BphS expression (**R11**. to **R15**.), 4) PA2133 expression (**R16** to **R19**), 5) Q expression (**R20** to **R24**) and 6) LKD expression (**R25** to **R30**). Details of these reactions are described below (Table S4-6):

**Photon activation of BphS.** Photon-activated diguanylate cyclase (DGC) BphS is stable as a dimer[14], which is denoted by Di_BphS. Using a pseudo-chemical reaction, the near-infrared Light (NIR) absorption of Di_BphS is modelled by reaction 1. R2 describes that the Di_BphS can convert from photon-absorption state (denote by Di_BphS_photon) to an activated state (Di_BphS$^{*}$) quickly, whereas slow relaxation of Di_BphS$^{*}$ to an inactivated state in absence of NIR-light (dark) is given by reaction 3. Note that Di_BphS$^{*}$ also can convert fastly to an inactivated state in presence of infrared light[14].

**c-di-GMP signaling.** Activated BphS enables the cyclization of two GTP molecules to form a c-di-GMP molecule (R4)[14]. R5 and R6 describe that phosphodiesterase (PDE) PA2133 can first bind to c-di-GMP molecule to enable hydrolysis of c-di-GMP in next[15]. FleQ from *Pseudomonas aeruginosa* is a c-di-GMP responsive transcriptional factor [16]. Either one or two c-di-GMP molecules can bind to one FleQ to form complexes (R7 and R8)[17]. R9 or R10 represent that the production or degradation c-di-GMP arose from the endogenous DGCs or PDEs existing in *P. aeruginosa[18].*

**BphS expression.** Constitutive promoter PA1/O4/O3 lead the coding sequence of *bphS* to form mRNA transcripts (R11), which carry the information for the subsequent translation, resulting the BphS synthesis (R12). R13, R14 or R15 describe the degradation/dilution of mRNA, inactivated BphS or activated BphS.

**PA2133 expression.** Constitutive promoter J23109 lead the coding sequence of *PA2133* to form mRNA transcripts (R16), which carry the information for the subsequent translation, resulting the PA2133 synthesis (R17). R18 or R19 describe the degradation/dilution of mRNA or PA2133.

**Q expression.** Complex formed by c-di-GMP and FleQ can bind to the c-di-GMP responsive promoter P*cdrA* to initialize the transcription (R20)[19]. Active promoter P*cdrA* lead the coding sequence of *Q* to form mRNA transcripts (R21), which carry the information for the subsequent translation, resulting the Q synthesis (R22). R23 or R24 describe the degradation/dilution of mRNA or Q.

**LKD expression.** Previous studies indicate that two Q promoters bind to a direct-repeat DNA site and contact distinct elements of the RNA exit channel, leading to the RNA polymerase resistant to termination signals (R25 and R26)[20]. Active promoter pR' lead the coding sequence of *LKD* to form mRNA transcripts (R28), which carry the information for the subsequent translation, resulting the LKD synthesis (R22). R29 or R30 describe the degradation/dilution of mRNA or LKD.

**2) Simulate and verify the computational model.**

Our main objective is to design synthetic genetic circuit to dynamic program bacterial lifestyles (planktonic, biofilm and lysis) via manipulating the LPD of NIR. We first obtained a set of parameter values (Table S6-7) that represents real behavioral data according to previous literature reports, and created pseudo input light signal (doses schedule are list in Table S9). Then, we used this model to simulate the behavior of each species in the experiment, such as the expression level of c-di-GMP and protein Q, and to investigate how species changes with different model parameters of experiments. We set the threshold of c-di-GMP for biofilm formation at 1 μM based on literature reports, and the threshold of LKD for bacteria lysis at 35 nM according to our previous experiment results. By adjusting the values of kinetic parameters within a reasonable range, we successfully simulated the result that the bacterial lifestyle can be dynamically programmed by manipulating the LPD of the NIR (Fig. S4 b). The value of the input photo signal in the model multiplied by a coefficient (10) is equivalent to the LPD ($mW/cm^2$) used in the experiment.

More importantly, we can change the kinetic parameters of a specific reaction in batches, and simulate its impact on the experimental results. This helps us quantitatively analyze and explore key parameters, and to develop the optimal experimental protocols for subsequent screening and construction of strains capable of switch lifestyle in response to illumination with NIR light of reasonable ranges of LPD (Fig. S4).

**3) Recalibrate the model parameters from experimental data.**

We first constructed a bacteria strain only containing NIR light-responsive module and c-di-GMP hydrolysis module. Through altering the expression of *PA2133* by replacing the sequence of promoter or RBS, we could determine the threshold of LPD for promoting bacteria biofilm formation, and modified parameters (Reactions R1-10) in the model until the simulation results agreed with the experimental results. After identifying the expression elements of *PA2133* (J23109-RBS010), we constructed a strain containing the lysis system and tested the corresponding LPD range inducing bacterial lysis to calibrate the key parameters of reactions R20-30. The information of engineered strains used to calibrate the model parameters and the corresponding experimental results is listed in Table S8.

**Supplementary Note 6: High-throughput screening of strains capable of switch lifestyle in response to NIR.**

**1) A portable device for wireless control of illumination.**

Construction of strains capable of switching lifestyle requires multiple rounds of genetic engineering. Specifically, a series of strains with different expression levels of *PA2133* and *Q* genes were first constructed to quantitatively characterize the LPD range corresponding to various lifestyles. According to the experimental results, strains with programmable lifestyle were screened by adjusting the RBS before *PA2133* and *Q* gene. In this study, we developed a high-throughput assay for rapid screening of NIR responsive strains with a reasonable range of LPD. Firstly, we designed a portable device to provide different LPDs of red light for 96-well plate（Fig. S7）. It can be easily controlled by an app on smartphone or MATLAB on PC, each well has an LED that can work separately. When the setting numerical value changed from 0 to 255, the corresponding LPDs can be changed from 0 to 717 $\mu W/cm^2$ (measured at 680 nm). For example, setting numerical value were 0, 8, 15, the corresponding LPDs were 0,15.1,106 $\mu W/cm^2$

**2) High-throughput assay for rough screening of NIR responsive strains.**

All bacterial strains to be tested were cultured overnight in LB medium at 37 °C, and diluted 100× to fresh LB medium until $OD_{600} \approx 0.6$. The resultant cells were loaded 100 μL into 96-well plate illuminated by a 96-well plate LED array for 2 h at room temperature. Each sample has four duplicates at each LPD. Then, we recorded bacterial aggregation in the plate with a camera and measured the $OD_{600}$ of the supernatant with a microplate reader. When the $OD_{600}$ of the supernatant was half that of the darkness group, the corresponding LPD value was defined as the critical LPD value corresponding to bacterial lysis (Fig. S10). This high-throughput assay can rapidly characterize the NIR susceptibility of engineered strains within 2 hours.

**3) Crystal violet assay to determine the threshold of LPD for biofilm formation**

Briefly, Exponentially-growing *Pseudomonas* was harvested and the cell pellet was then resuspended in the fresh FAB medium based on desired concentration ($OD_{600}$ ≈ 0.5). Pipet 100 μL of each diluted culture into each well in a fresh microtiter plate. Then microtiter plate was exposed to various LPD of NIR illumination for 24 h at room temperature. After exposure, the bacterial liquid was gently transferred to another new 96-well plate and read absorbance at 600 nm by microplate reader. The initial 96-well plate was washed twice with PBS (200 μL) before staining with crystal violet (125 μL; 0.1% crystal violet in PBS). After 10 min, the excess crystal violet was removed and plates were washed twice and air dried. Biofilms that had formed were visible at the bottoms of 96 black wells. The amount of remaining crystal violet stain in each biofilm was quantified after addition of 33% glacial acetic acid (125 μL) followed by mixing and measurement of the $OD_{570}$ using microplate reader.

**Supplementary Note 7: Measurement of NIR LPD for manipulating bacterial lifestyle.**

To study the bacterial lifestyle transitions over time, we used microscope for accurate control of LPD of NIR, and recorded the number of surface-adhering bacteria in situ under different illumination conditions. To determine the LPD of NIR (680 nm) illuminated on bacteria, we measured the power at outlet of the 60× oil objective (Olympus) using a power meter (Newport 842-PE) and converted by reduction of the exposure time and the area of one exposed microscope filed. The illumination scheme set the parameter for exposure time, exposure strength, light source and shooting frequency.

In the mouse experiment, the LPD (measured at 680 nm) of red light in the cage was adjusted by changing the number of LED lamp beads that emit red light around the cage. The red LED strip lights were fixed outside the cage with double-sided tape and connected to the power socket through a 45W transformer (DC12V). One-meter-long light strip could just make a full circle around the cage, and there were 60 LED light beads in it. We could control the illumination time and the LPD of red light in the cage by controlling the switch of the timing power socket meter. Each socket controlled a transformer that connects LED strips of different lengths.

**Supplementary Note 8: Ex vivo assays for tumor tissues.**

**1) Hematoxylin and eosin (H&E) and TUNEL staining of tissue sections for the subcutaneous solid tumor.**

After a sequence of treatments with engineered bacteria strains or PBS under different illumination schemes, tumor tissues were harvested, fixed and embedded into a Tissue-Tek OCT compound (Sakura Finetek, Torrance, CA). Then, the tissues were sectioned into 8-μm-thick slices with a -20 °C Leica CM1950 cryostat, and transferred to glass slides. Tissue slices were processed with H&E Staining Kit (Beyotime) as directed by the manufacturer's instructions. Subsequently, the slices were mounted with anti-Fade Medium (Sangon) and imaged using ZEISS Advanced Upright Microscope and TissueFAXS PLUS (TissueGnostics Gmbh).

TUNEL (Beyotime) was used to analyze the apoptosis in the tumor sections. The nucleus was stained with DAPI (Sangon). The operation procedure of the staining experiment was carried out according to the manufacturer's protocol. Fluorescence images was acquired by a spinning-disc confocal microscope (IX81, Olympus). Apoptotic cells stained by TUNEL was green, while the nucleus was blue.

**2) IVIS Spectrum images of solid tumors**

Due to the limitation of our experimental conditions, we could not detect the fluorescence intensity of living mice in situ with IVIS Spectrum (Perkin Elmer), but only monitored the fluorescence intensity of fixed tumor tissues. H017 was labelled with GFP fluorescent protein, so the changes in the number of bacteria within the tumors could be obtained by monitoring the changes in fluorescence intensity over time.

Tumor growth was tracked daily by monitoring tumor volume. A549 tumor-bearing mice intratumorally injected with H017 were treated with different illumination schemes, the tumor tissues were isolated and fixed with 4% paraformaldehyde overnight at 4 °C. Fluorescence images of tumor tissues were then acquired using IVIS and analyzed by Living image version 4.5.2 software (PE).

**3) Characterization of lytic stability of intratumoral bacteria.**

To test whether intratumoral bacteria H017 can be lysed, tumor tissues irradiated with Middle-LPD NIR for 7 days were collected, weighed, homogenized, serially diluted, and plated on LB agar plates with or without gentamicin. We picked 96 single colonies and inoculated them into fresh LB medium (200 μL) in a 96-well plate. Then, 100 μL bacterial culture was transferred into another 96-well plates and wrapped with tin foil. Subsequently, these two 96-well plates were placed in a shaker irradiated with red light for 8 hours at 37 °C. Finally, the absorbance at 600 nm of bacterial culture in 96-well plates was measured with a microplate reader to evaluate the growth of bacteria.

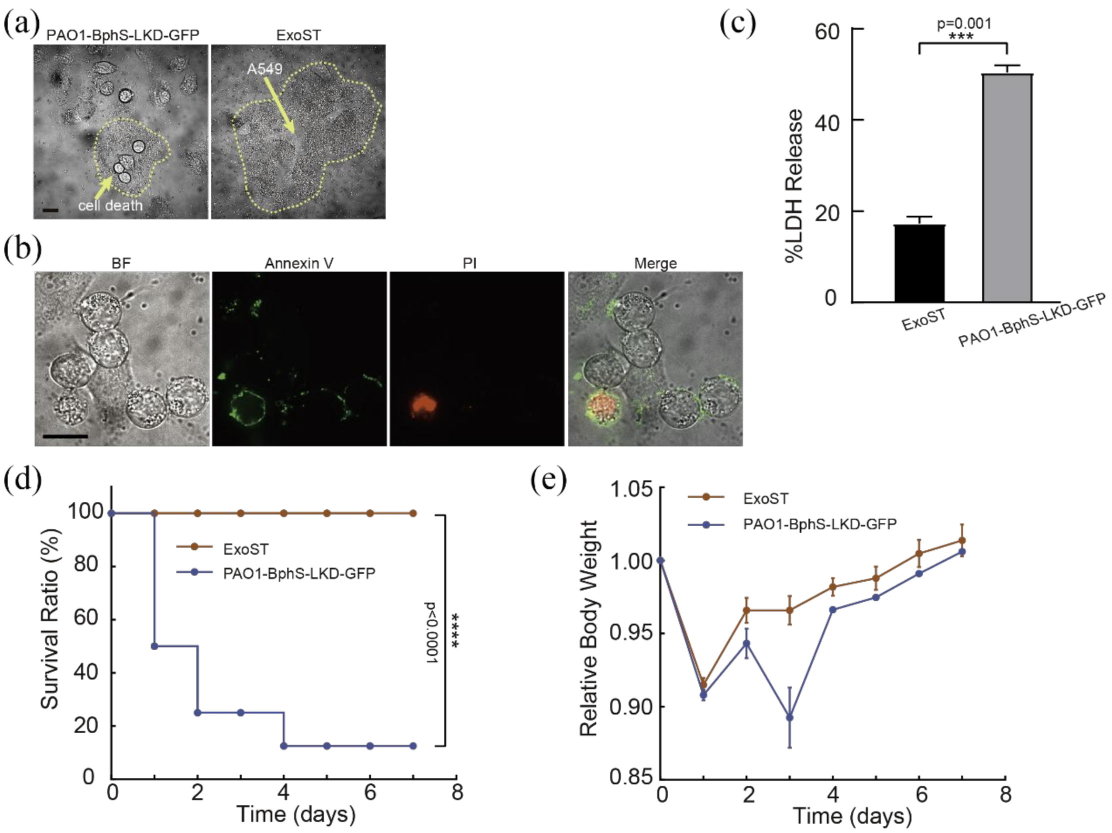


**Figure. S1. Attenuation of *P. aeruginosa*.** (a) Representative BF images of A549 cells co-cultured with PAO1-BphS-LKD-GFP or ExoST in microfluidic devices under middle-LPD NIR illumination for 5 h and 8h respectively. (b) Representative BF, Annexin V staining (green), PI staining (red) and merge images of A549 cells co-cultured with PAO1-BphS-LKD-GFP under darkness for 5 h. (c)LDH release assay of A549 cells co-cultured with PAO1-BphS-LKD-GFP or ExoST. A549 cell alone was used as control. (d) Survival ratio and (e) relative body weight of mice. A549-bearing mice (n=8 per group) were challenged with s.c. injection of $5\times10^{7}$ CFU ExoST or PAO1-BphS-LKD-GFP at day 0 (***P<0.001, unpaired two-tailed t-test). Error bars represent SEM. Scale bars for all images are 20 μm.

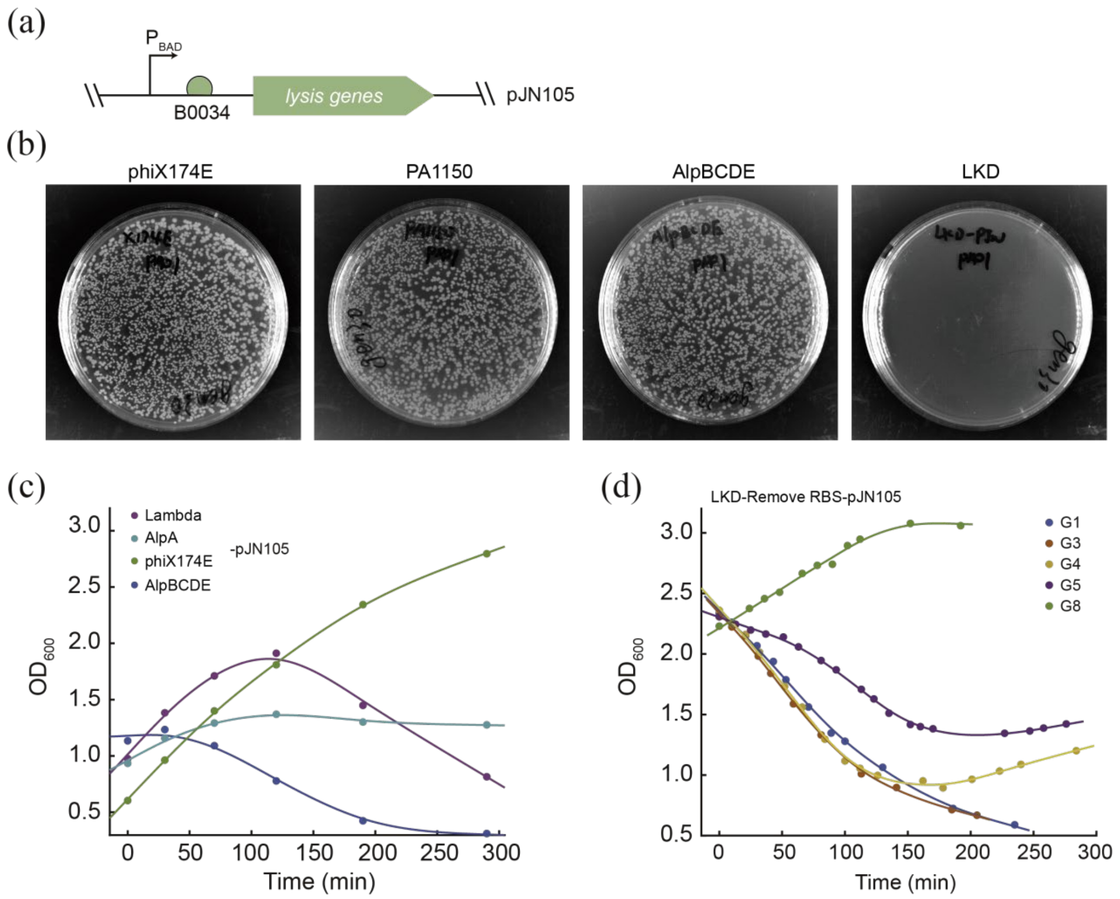


**Figure. S2. Screening for lysis genes that can induce lysis in *P. aeruginosa*.** (a) Schematic diagram of plasmid used for sorting suitable lysis genes. (b) The digital photographs of agar plates after electroporation of various plasmids and incubated at 37°C for 12 h. (c) Representative growth curves of bacterial strains containing different lysis genes after supplemented with 0.2% arabinose. (d) Growth curves of bacterial strains containing plasmid *LKD*-Remove RBS-pJN105 after supplemented with 0.2% L-arabinose, different generations were tested.

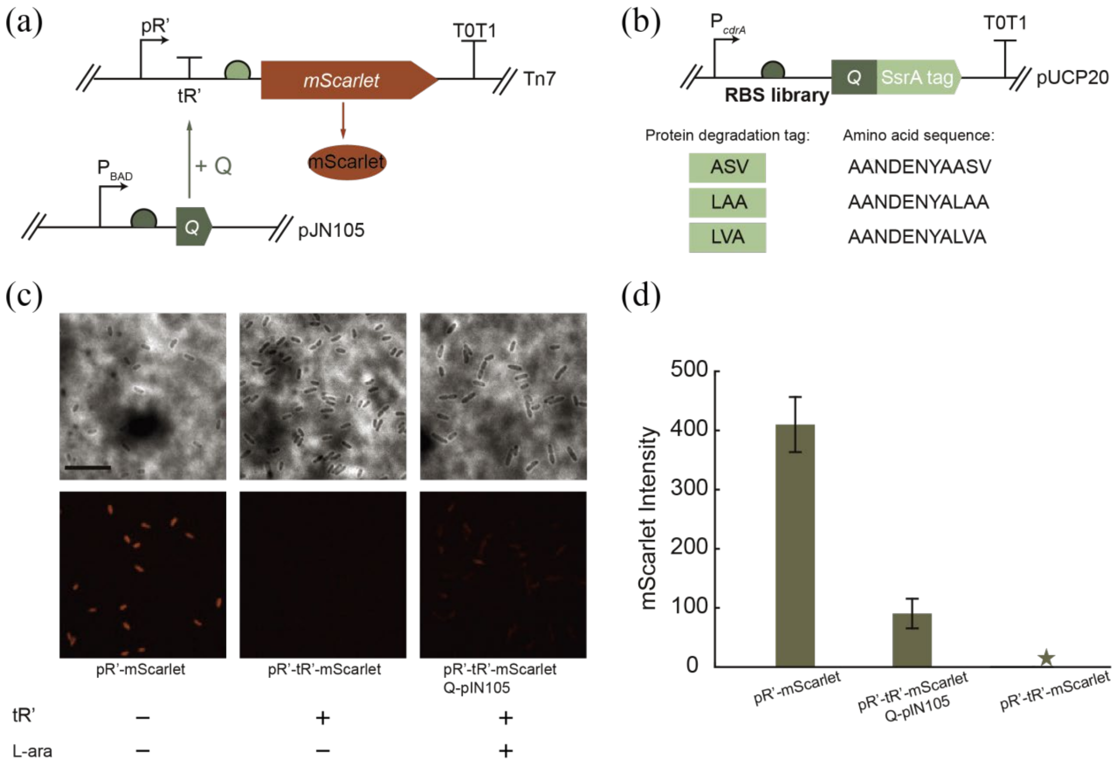


**Figure. S3. Minimize the leaky expression level of lysis cassette.** (a) Schematic diagram of plasmid used for testing the effect of reducing leaky gene expression by pR'-tR' late expression system originated from λ phage. (b) Schematic diagram of key sequence elements to be optimized, including RBS and ssrA protein degradation tag. (c)Representative BF and RFP images of different bacteria strain. 0.2% L-arabinose was supplemented for expression of Q. Scale bar, 10 μm. (d) Fluorescence intensity measured in (c). The star means no value detected. Error bars represent SEM.

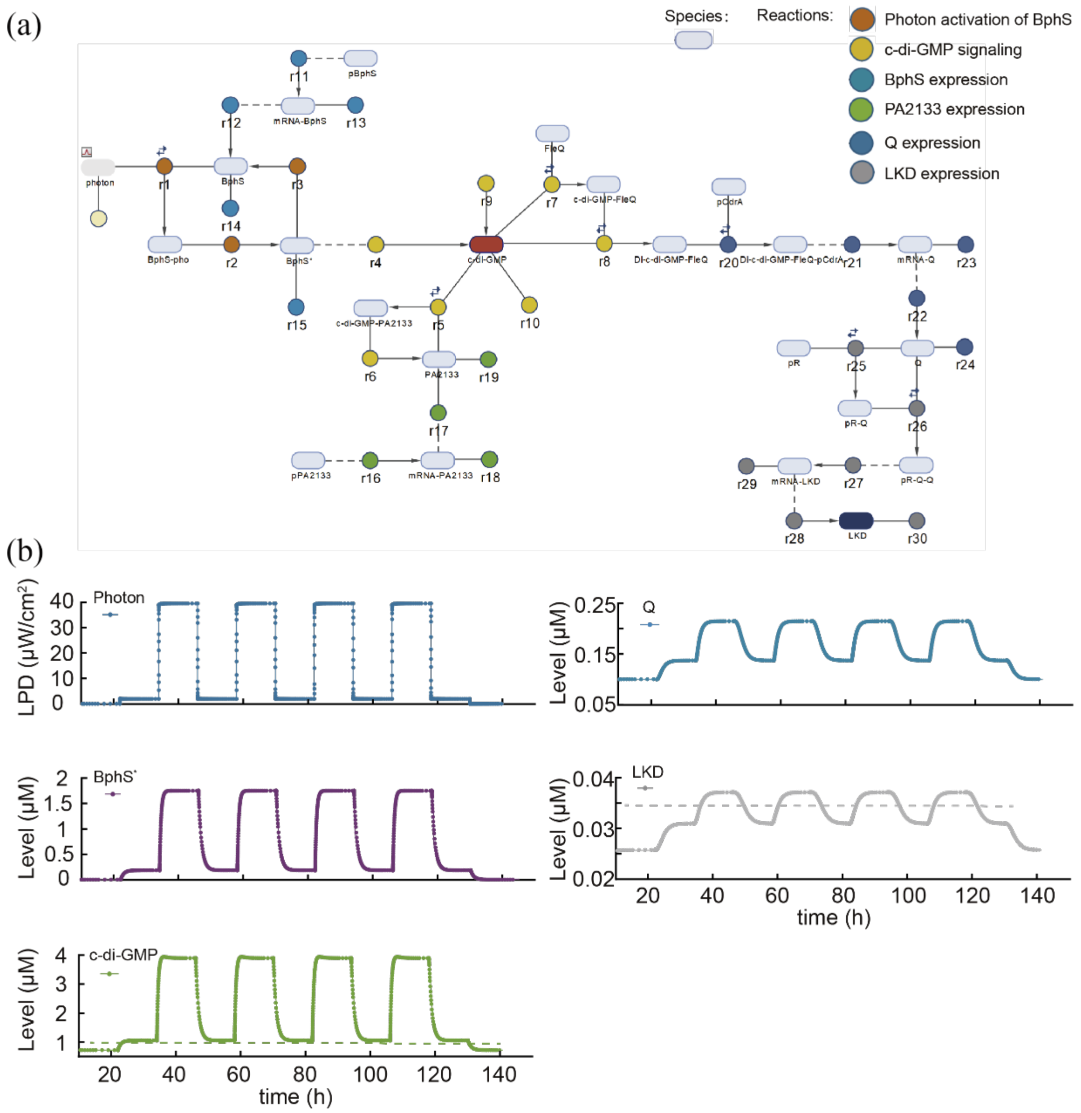


**Figure. S4. Modeling of dynamic programming bacterial lifestyle transition.** (a) System biology model of gene regulation pathway. The ovals represent the species, and the circles represent reactions in the model. Each reaction was numbered, denoted as r1-r30. (b) The key species concentrations over time were simulated after the input photon signal was created.

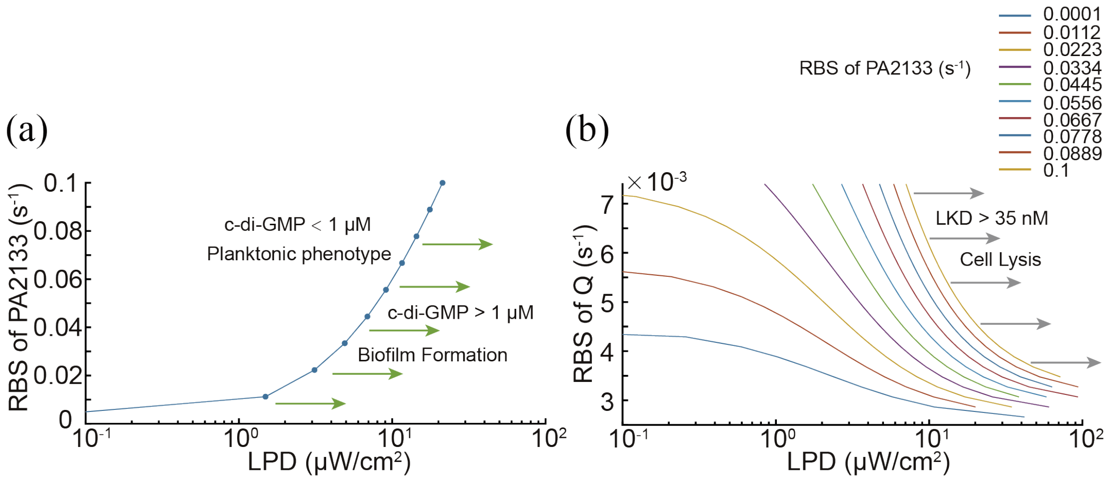


**Figure. S5. Simulation results for the different parameter changes.** (a) The simulation result shows the relationship between the threshold of LPD for biofilm formation and the intensity of RBS in front of PA2133. (b) The RBS in front of PA2133 was settled when simulating the effect of Q gene expression level on the threshold of LPD for bacterial lysis.

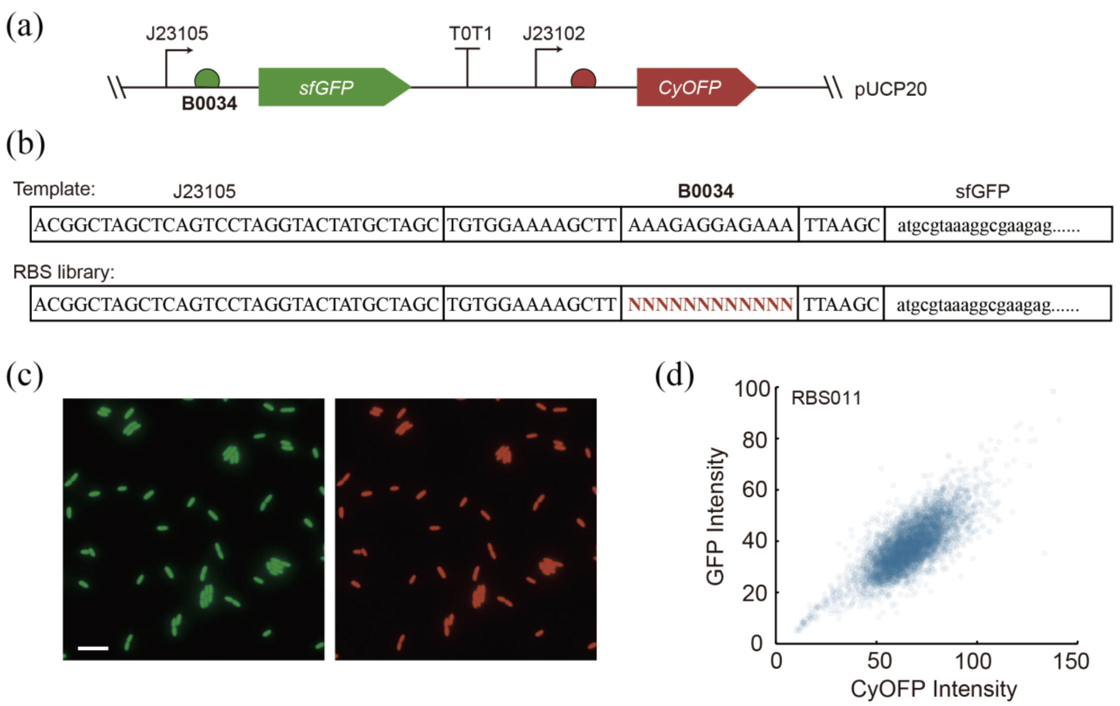


**Figure. S6. Construction and characterization of RBS library.** (a) Schematic diagram of plasmid used for quantifying RBS strength and (b) base sequence of RBS library. (c) Representative fluorescence images of GFP and CyOFP obtained from RBS011. (d) Fluorescence intensities measured at single-cell level of multiple cells (>5000) with RBS011. Each dot indicates one bacterium counted. Scale bar, 10 μm.

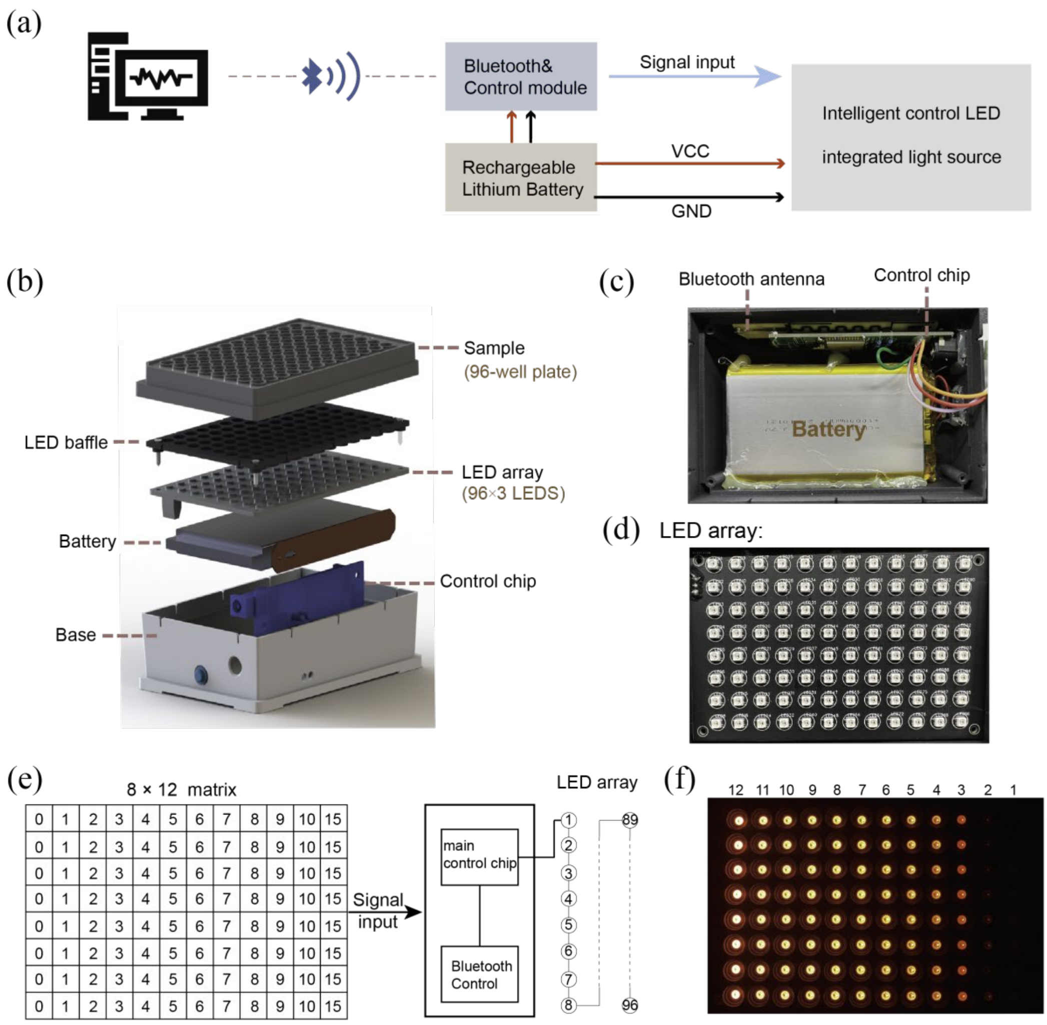


**Figure. S7. Design of a portable device for wireless control of illumination.** (a) Schematic diagram of the working principle of the device. (b) Schematic diagram of the device structure. The rechargeable lithium battery provides energy for the whole device. The signal input is provided by PC or mobile phone through Bluetooth, LED array is controlled by the control chip. The assembling graph of the device and physical maps of key parts include (c) Bluetooth antenna, control chip, battery and (d) LED array. (e) The input signal is generated by an 8×12 matrix corresponding to 96-well plate. Each LED can be set an integer value from 0 to 255 separately, the lager value corresponds to higher LPD. (f) The digital photograph of the device when set the value matrix shown in (e) by PC.

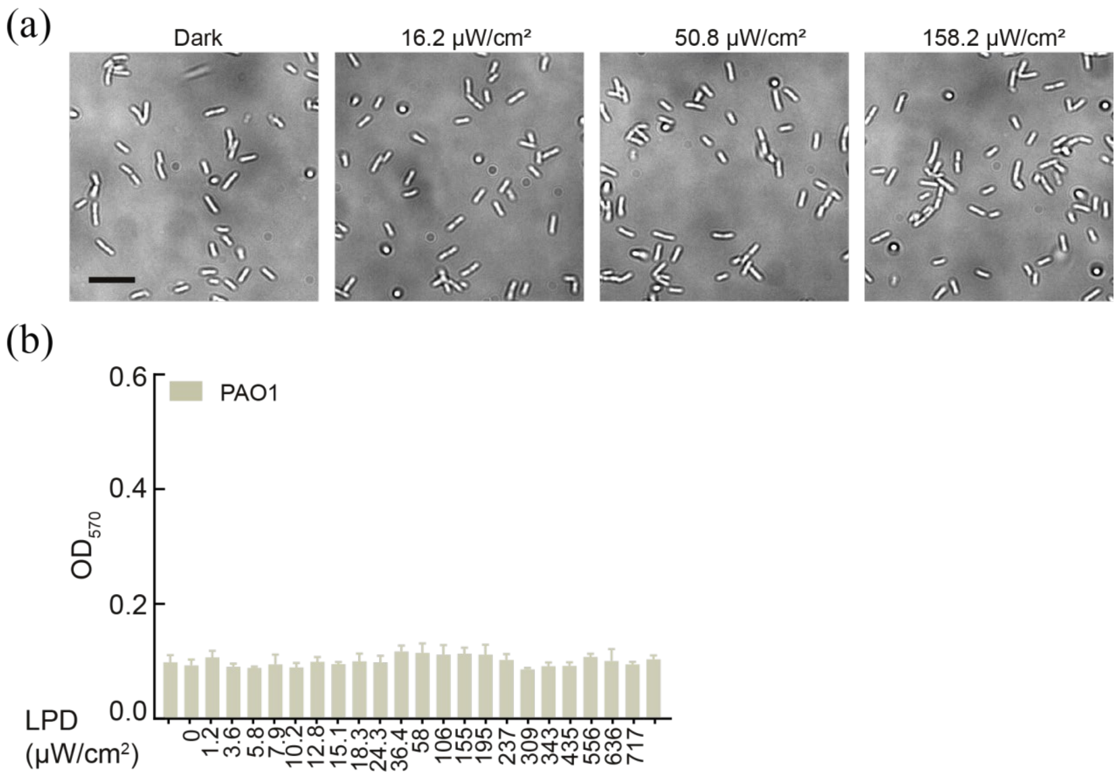


**Figure. S8. Wild type PAO1 cannot respond to NIR.** (a) Representative BF images of PAO1 wild type illuminated with different LPDs for 8 h in microfluidic devices. Scale bar, 10 μm. (b) PAO1 was cultured statically in a 96-well plate illuminating with different LPDs for 24 hours. The resultant measurement of $OD_{570}$ after crystal violet staining (n = 4 biological replicates per group). Error bars represent SEM.

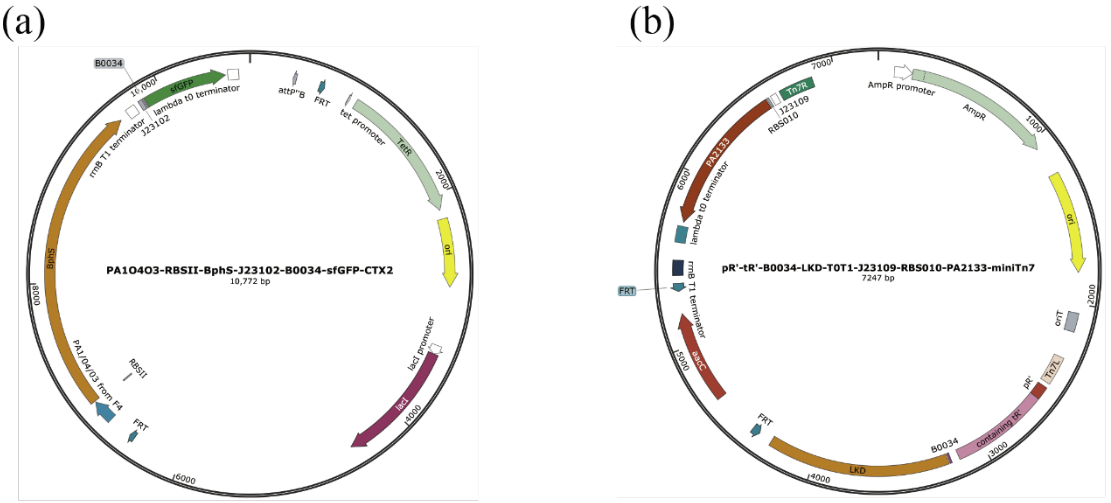


**Figure. S9. Plasmid map for insertion of functional modules into the PAO1 genome.** (a) Plasmid used to integrate the NIR light responsive module BphS and fluorescent protein GFP into the chromosomal *attB* site. (b) The c-di-GMP hydrolysis module PA2133 and lysis cassette LKD was inserted into the genome *attTn7* site using miniTn7 plasmid.

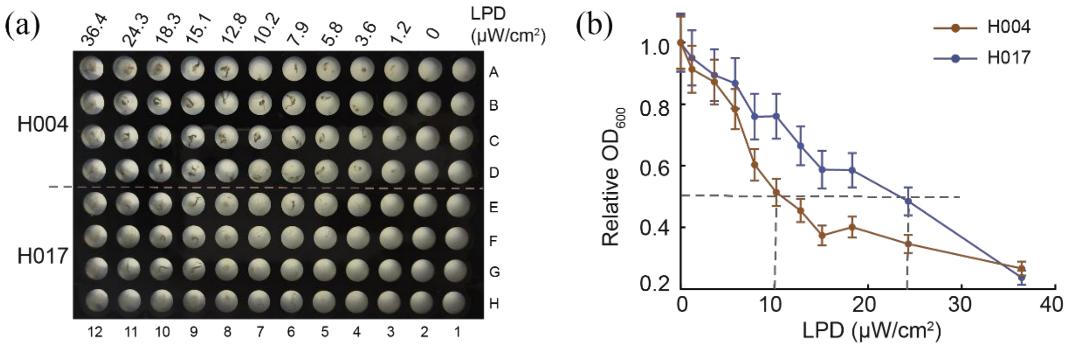


**Figure. S10. Roughly screen bacterial strains through 96-well illumination device.** (a) Overview of H004 and H017 treated with different LPD illumination for 2 hours in 96-well plate. (b) Relative $OD_{600}$ of supernatant in (a) was measured by microplate reader.

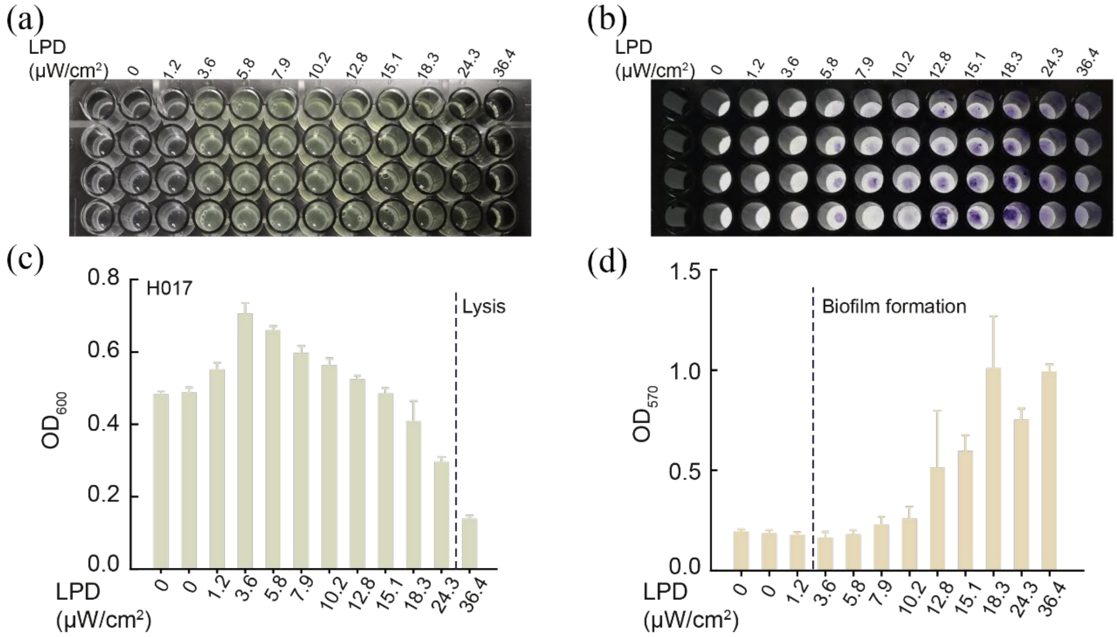


**Figure. S11. Characterization of bacterial lysis and biofilm formation.** (a) Overview of H017 in 96-well plate treated with different LPD for 24 h and (c) measurement of $OD_{600}$. (b) Crystal violet staining results of H017 and (d) measurement of $OD_{570}$. Error bars represent SEM, n = 4 biological replicates per group.

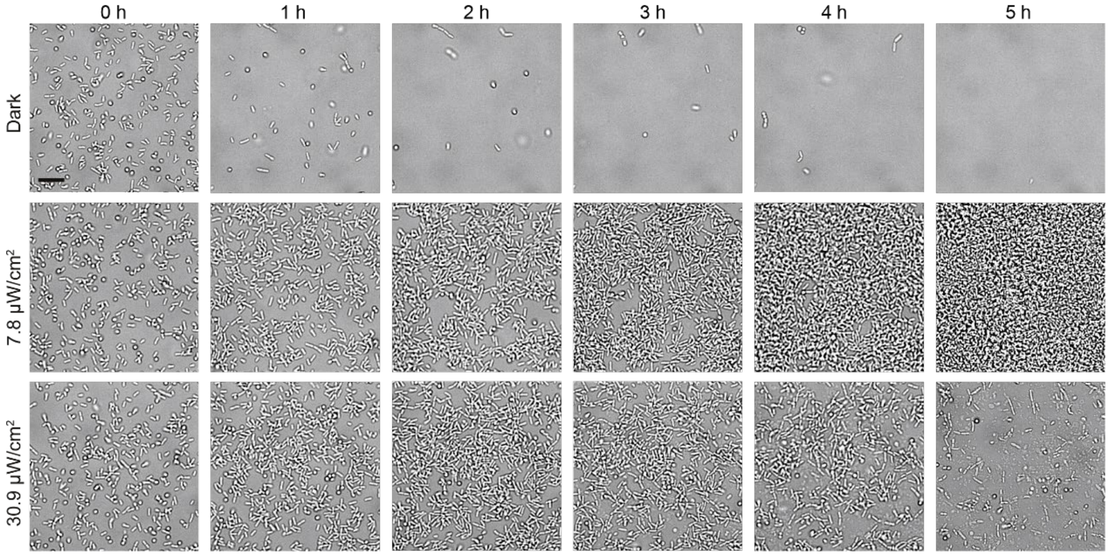


**Figure. S12. Time-lapse BF images of H017 illuminated with different LPDs of NIR.** BF images of H017 illuminated with different LPDs for 5 hours in microfluidic devices. Images were acquired at 30-min intervals. Scale bar, 10 μm.

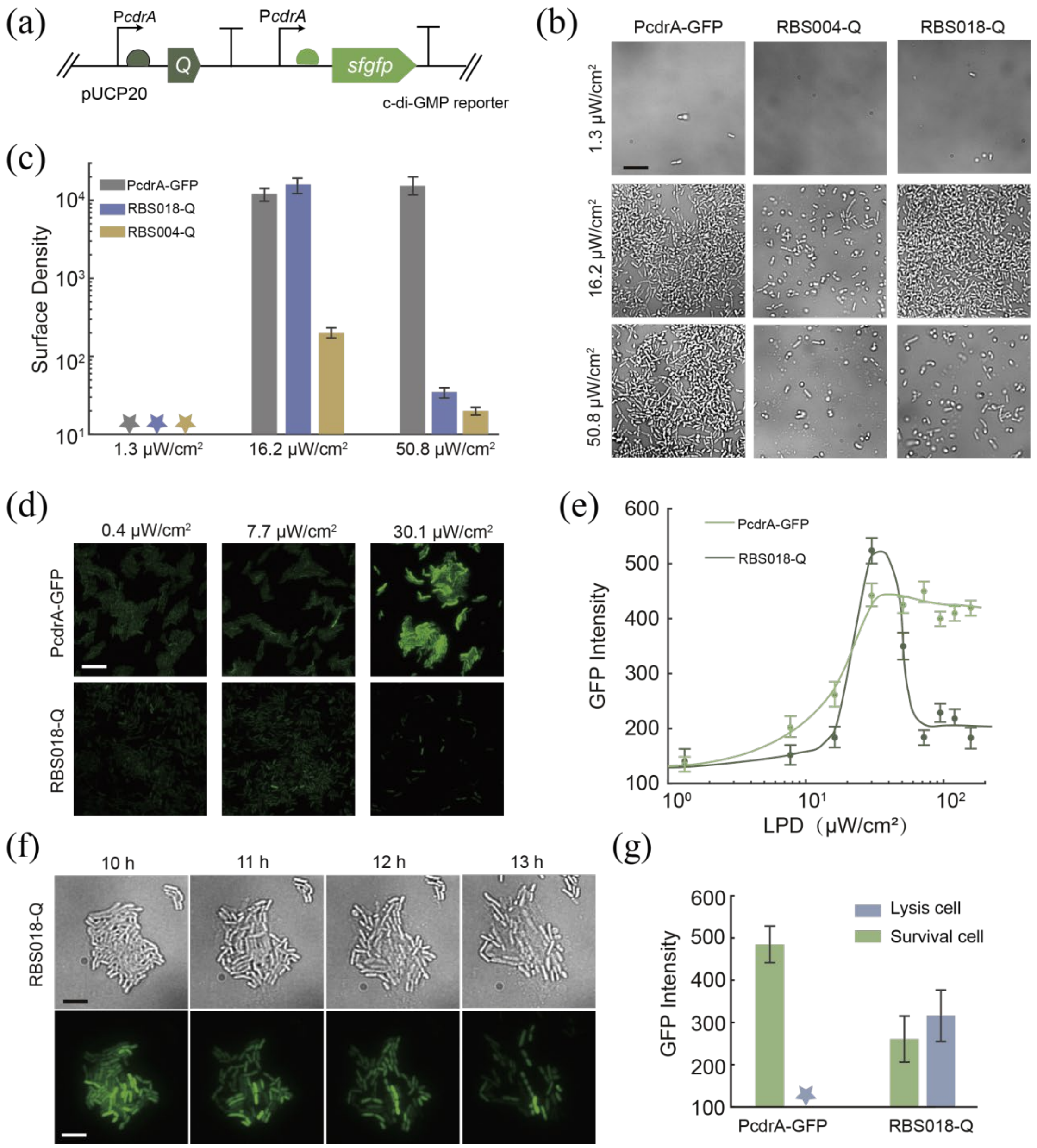


**Figure. S13. ExoST harboring varied plasmids show different LPD responsive properties.** (a) Schematic illustration of plasmid used for detection c-di-GMP levels in lysing bacteria. (b) Representative BF images of PcdrA-GFP, RBS004-Q and RBS018-Q after irradiated with different LPD of NIR for 12 h, and (c) surface density was calculated at last. (d) Representative GFP images of PcdrA-GFP and RBS018-Q illuminated with different LPD for 16 hours. (e) Nine LPDs were selected and GFP intensities were analyzed separately after 8 h illumination. (f)Time-lapse BF and GFP images of RBS018-Q under high-LPD illumination (30.1 μW/cm$^2$). (g) GFP intensities of lysed cells and viable cells of PcdrA-GFP and RBS018-Q measured after NIR illumination for 16 h. The star means no value detected. Scale bars are 10 μm for all images.

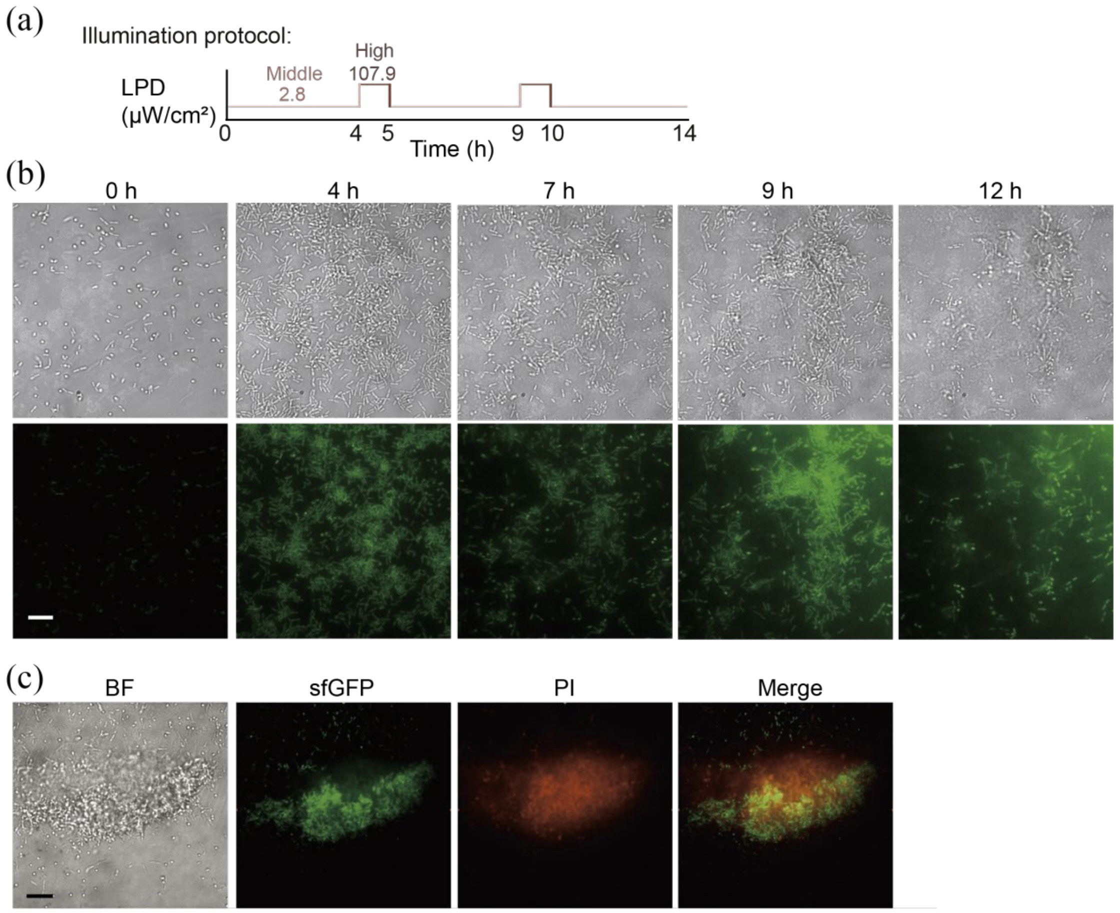


**Figure. S14. Two biofilm-lysis lifestyle transition in vitro.** (a)Schematic diagram of illumination scheme applied. (b) Time-lapse BF and GFP images of H017 treated with illumination scheme in (a). (c) BF, sfGFP, PI, GFP&PI merge image of H017 after 7 hours of illumination. Live bacteria were indicated by GFP and dead bacteria were marked by PI. Scale bars for all images are 10 μm.

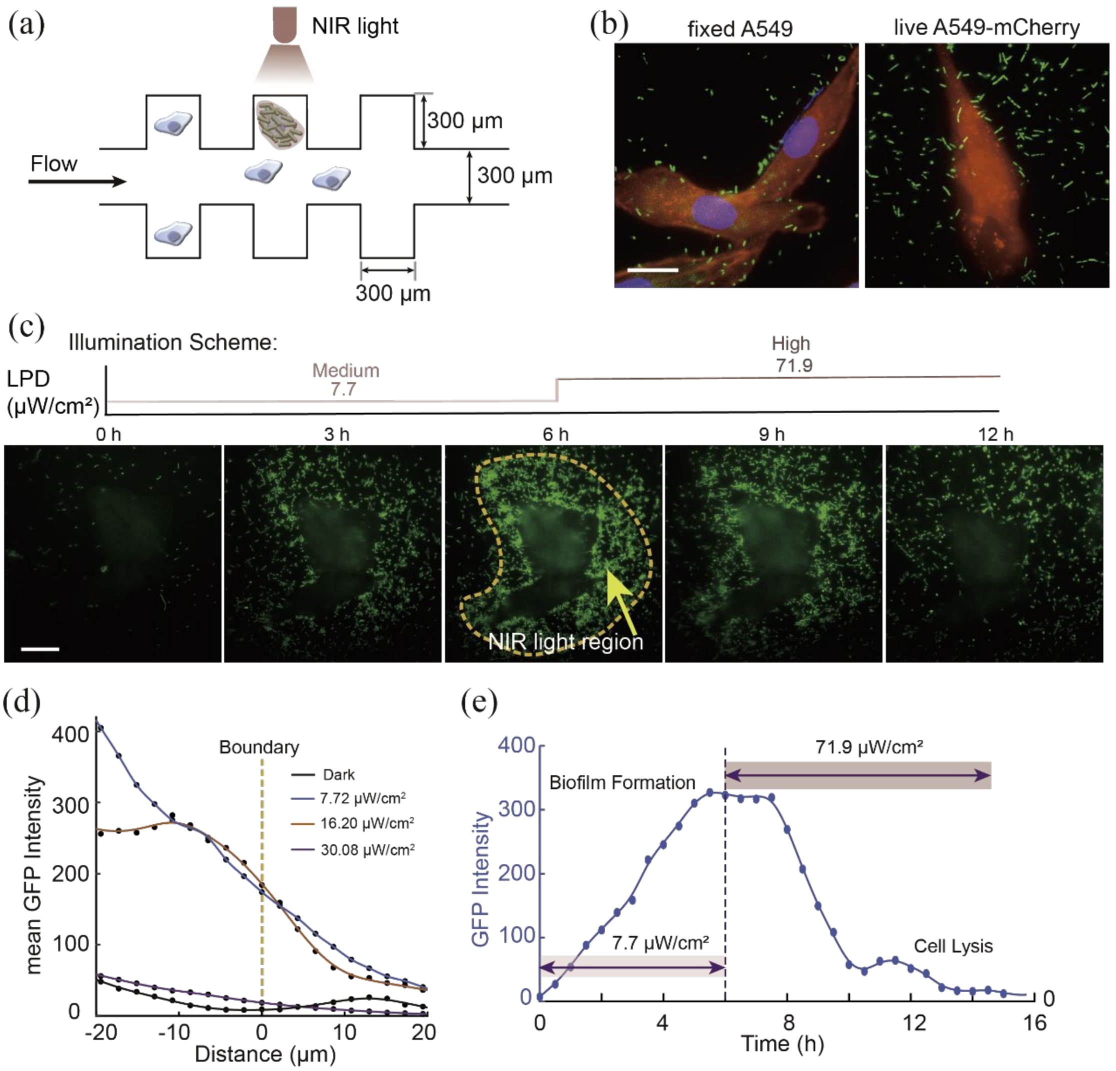


**Figure. S15. Manipulation of bacterial lifestyles at single cell resolution in vitro**. (a) Schematic illustration of the microfluidic device for bacterial cell co-culture. (b) Representative fluorescence images of the initial co-culture of fixed A549 or live A549-mCherry with RecA-H036. (c) Time-lapse GFP images of H017 treated with middle LPD for 6 h and then high-LPD for 6 h. The boundary of NIR illumination region is indicated by yellow dashed lines. (d) Mean GFP intensity distribution with distance from the illumination boundary under treatment with different LPDs for 8 h. Distance here was defined that inside the boundary is negative and outside the boundary is positive. (e) Mean GFP intensity in (c). Scale bars are 20 μm for all images.

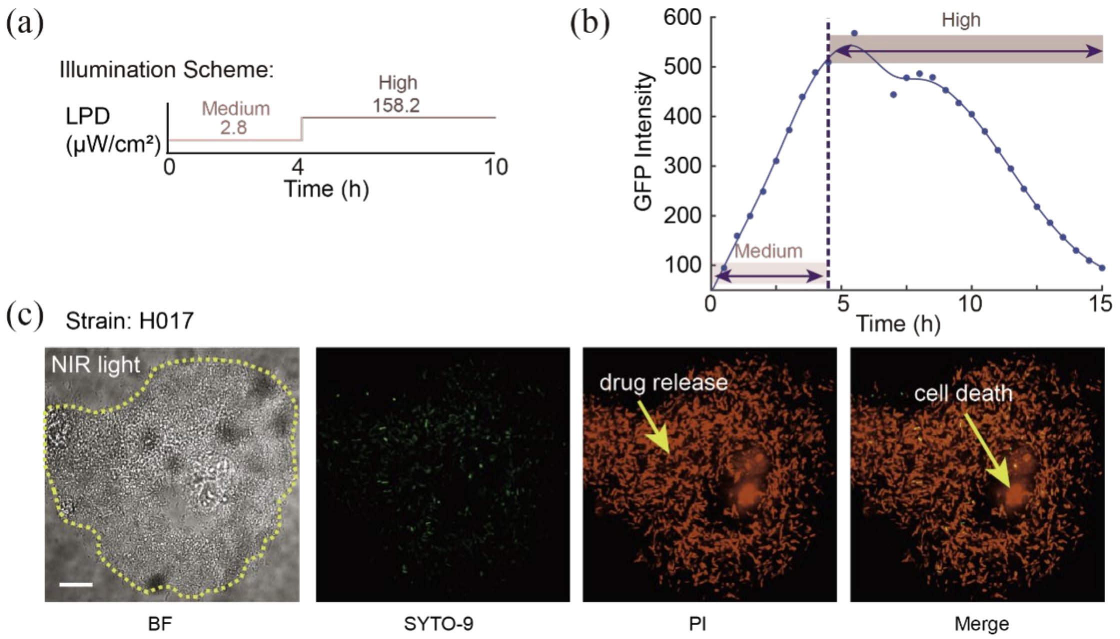


**Figure. S16. Biofilm-lysis lifestyle transition results in the death of live A549 cells.** (a)Schematic diagram of illumination scheme applied. (b) Mean GFP intensity of H017 treated with scheme in (a). (c)Representative BF, H017 (green), PI staining (red) and merge images of A549 and H017 after co-culturing for 10 h. Scale bar, 10 μm.

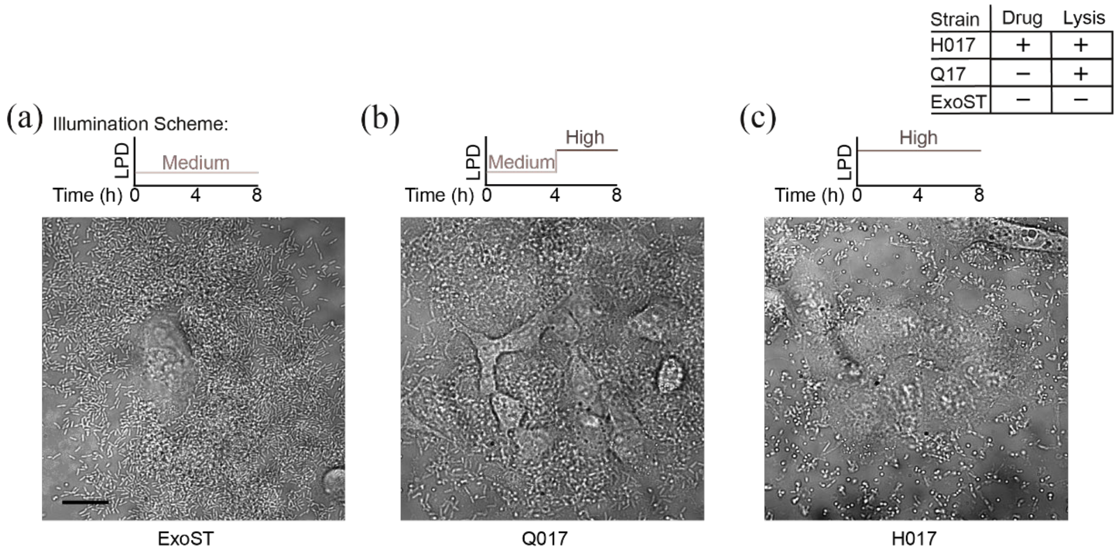


**Figure. S17. Bacterial strains co-cultured with A549 live cells and treated with different illumination schemes.** Representative BF images of (a) ExoST, (b) Q017, and (c) H017 co-cultured with A549 in microfluidic devices for 8 h, corresponding illumination schemes were shown above the images. Scale bar, 20 μm.

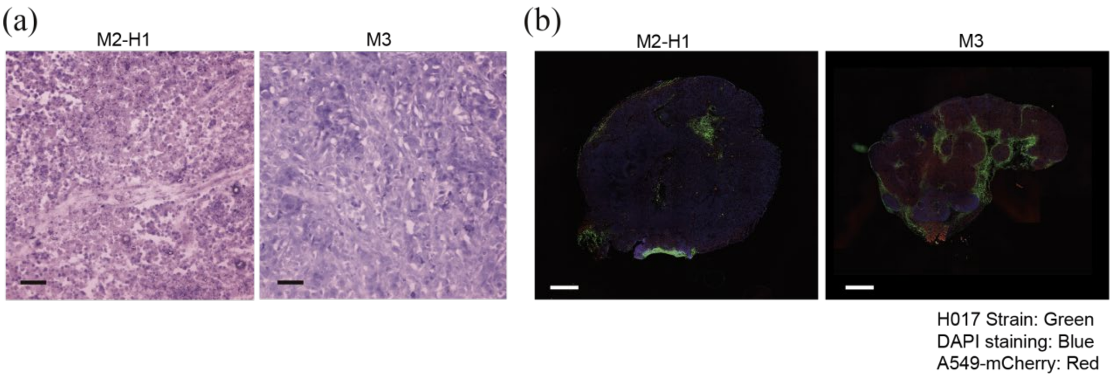


**Figure. S18. Histological analysis and panoramic fluorescence images of tumor sections treated with different NIR illumination schemes.** A549-mCherry-tumor-bearing mice received i.t. injection of 5×$10^7$ CFU H017 and treated with M2-H1 or M3, tumors were isolated at day 3. (a) H&E staining results of tumor sections. Scale bar, 50 μm. (b) Merged fluorescence images of tumor sections.H017 is green, A549-mCherry is red and DAPI is blue. Scale bar,1 mm.

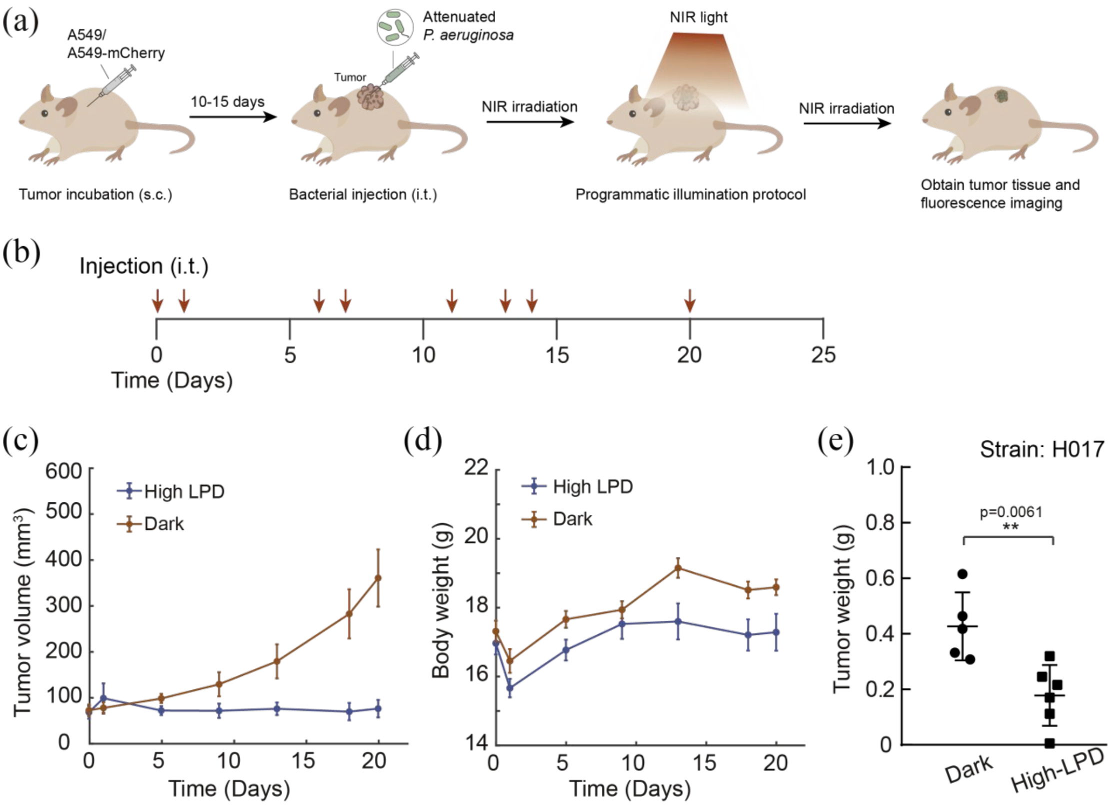


**Figure. S19. Multiple intratumoral injections of H017 and illumination of high LPD repress tumor growth.** A549-mcherry tumor cells were subcutaneously injected at day 0. When tumor volume reached about 100 $mm^3$, mice were divided into two treatment groups: illuminated with high-LPD NIR (n = 6) or kept in dark (n = 5) continuously. The tumors were finally isolated and used for immunostaining. (a) Experimental workflow of tumor-bearing mice treated with NIR light. (b) Mice were treated with i.t. injections of $5\times10^7$ CFUs H017 at the indicated time points (red arrow). (c) Changes in tumor volume and (d) body weight. (e) Distribution of tumor weight at day 25 (**P<0.01, unpaired two-tailed *t*-test). Data are representative of two independent experiments. Error bars represent SEM. Each circle or square represents an individual animal.

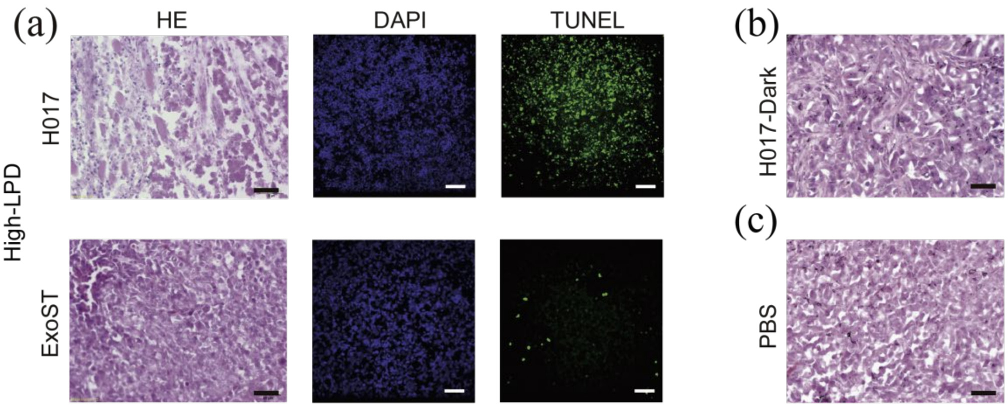


**Figure. S20. Histological analysis of tumor sections.** (a) Representative HE, DAPI and TUNEL staining results of tumor sections isolated from mice injected with H017 or ExoST and treated with high-LPD NIR. (b) Representative HE staining results of tumor sections isolated from mice injected with H017 and grew in dark conditions or PBS and incubated normally. Scale bar, 50 μm.

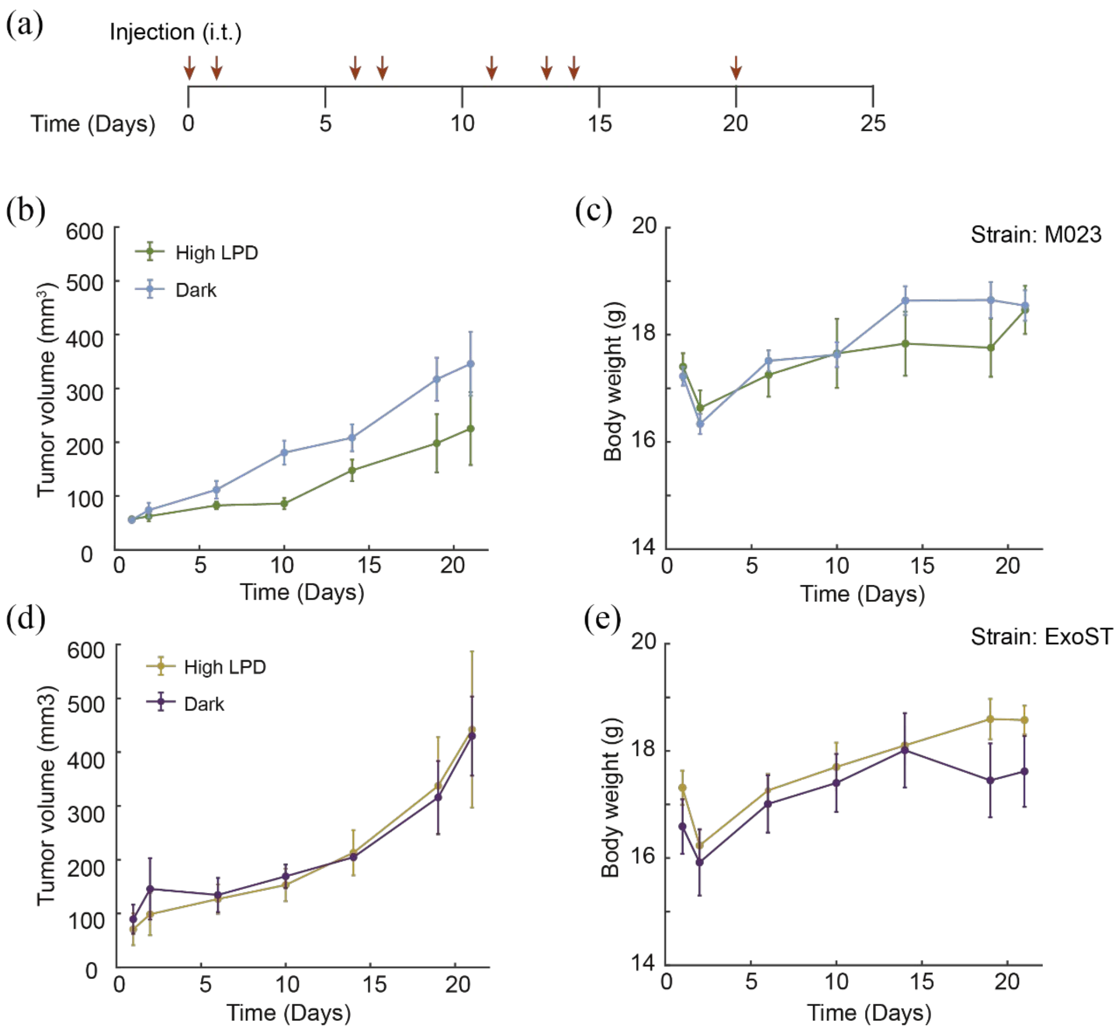


**Figure. S21. Effects of multiple intratumoral injections of M023 and ExoST on bacterial tumor growth under high-LPD illumination or darkness.** (a) A549-mcherry tumor bearing mice (n = 5 per group) were treated with i.t. injections of $5\times10^7$ CFUs engineered bacteria at the indicated time points (red arrow). (b, d) Changes in tumor volume and (c, e) body weight of mice treated with M023 and ExoST respectively. Error bars represent SEM.

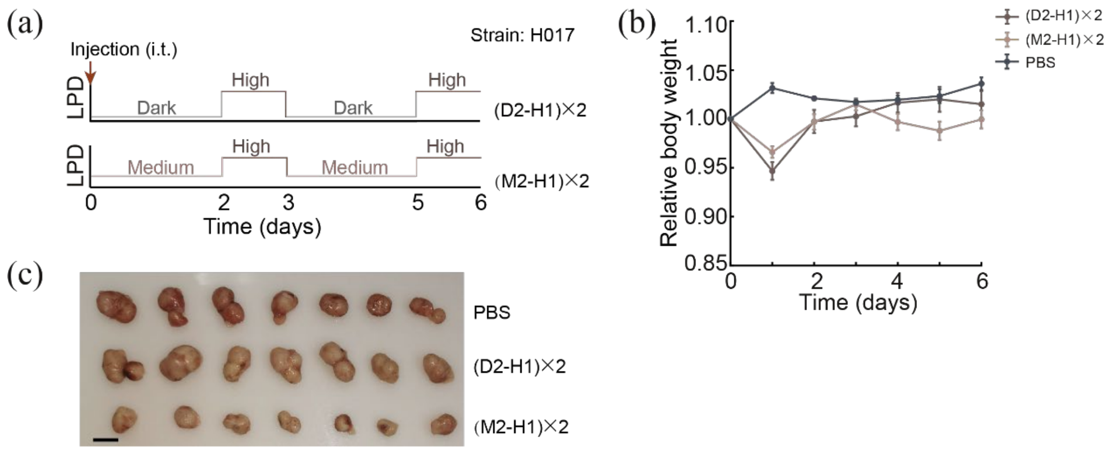


**Figure. S22. Biofilm-lysis lifestyle transition represses tumor growth in vivo.** (a) Schematic diagram of NIR illumination scheme applied. (b) Relative body weight changes of A549-mCherry bearing mice (n = 7 per group) treated with different illumination schemes. (c) The digital photographs of tumors isolated from each group of mice. Error bars represent SEM. Scale bar,1 mm.

#### Table S1. Attenuated *Pseudomonas* preferentially accumulates in the lung.

| Strains | Mice | Injection Method | CFU | Survial rate | Accumulate site | References |
|---|---|---|---|---|---|---|
| PAO1 | C57BL/6 | intratracheal | $5 \times 10^6$ | 0% | Lung | [21] |
| PAO1-ΔbphO | C57BL/6 | intratracheal | $5 \times 10^6$ | 90% | Lung | [21] |
| PAO1-ΔpvdQ | C57BL/6 | intratracheal | $5 \times 10^6$ | 70% | Lung | [21] |
| PAO1 | C57BL/6 | intranasal | $7.7 \times 10^5$ | 100% | Lung | [22] |
| PAO1 | C57BL/6 | intranasal | $1\times 10^7$ | n.d. | Lung/Spleen/Blood | [23] |
| PAO1-ΔpopB | C57BL/6 | intranasal | $1\times 10^7$ | n.d. | Lung | [23] |
| PAO1-ΔexoST | C57BL/6 | intranasal | $1\times 10^7$ | n.d. | Lung | [23] |
| PA14 | C57BL/6 | intranasal | $2.5 \times 10^5$ | 5% | Lung | [24] |
| PA14-ΔpvdF | C57BL/6 | intranasal | $2.5 \times 10^5$ | 70% | Lung | [24] |

#### Table S2. The colonization of *Pseudomonas* strains in tumor.

| Strains | Mice | Injection Method | CFU | Tumor | Antitumor effect | preferentially accumulate site | References |
|---|---|---|---|---|---|---|---|
| PA14 | BALB/c | intratracheal | $1\times 10^7$ | CT26 | n.d. | Tumor/Liver | [25] |
| PA14-ΔPA0781 | BALB/c | intratracheal | $1\times 10^7$ | CT26 | n.d. | Tumor (Liver none) | [25] |
| PAO1/PA14/ *P. putida* KT2440 | BALB/c | intravenous | $5 \times 10^6$ | CT26 | n.d. | Tumor | [26] |
| PA14 | BALB/c | intravenous | $5 \times 10^6$ | CT26 | + | Tumor | [27] |
| PA14 | BALB/c | intravenous | $5 \times 10^6$ | CT26 | + | Tumor | [28] |

#### Table S3. Plasmids used to screen lysis genes in *P. aeruginosa*.

| Plasmid | Relevant characteristics | Isolates |
|---|---|---|
| *phiX174E*-pJN105 | pJN105 inducible vector with fragment of *phiX174E*, Lysis protein E from Enterobacteria phage phiX174 | Yes |
| *PA0985*-pJN105 | S-type pyocin | Yes |
| *PA1150*-pJN105 | S-type pyocin | Yes |
| *PA3866*-pJN105 | S-type pyocin | Yes |
| *PA0614-PA0629*-pJN105 | the holin (*PA0614*) and the endolysin (*PA0629*) | Yes |
| *prtN*-pJN105 | a positive regulator of pyocin biosynthesis gene | Yes |
| *alpA*-pJN105 | lysis phenotype activator | Yes |
| *alpBCDE*-pJN105 | lysis cassette from PAO1 | Yes |
| *Lamda*-pJN105 | Lysis genes from phage λ | Yes |
| *LKD*-pJN105 | Lysis genes from *Pseudomonas* specific phage LKD16 | None |
| *LKD*-Remove RBS-pJN105 | Remove RBS upstream of *LKD* | Yes |

#### Table S4. Chemical reactions for the model.

| # | Reaction Equation | Description | Category |
|---|---|---|---|
| R1 | $Di\_BphS + photon \xleftrightarrow{k_{1r},\ k_{1f}} Di\_BphS\_photon$ | NIR-light absorption of BphS dimer | Photon activation of Bphs |
| R2 | $Di\_BphS\_photon \xrightarrow{k_2} Di\_BphS^*$ | Activation of BphS dimer | |
| R3 | $Di\_BphS^* \xrightarrow{k_3} Di\_BphS$ | Deactivation of BphS dimer | |
| R4 | $Di\_BphS^* \xrightarrow{k_4} Di\_BphS^* + cdiGMP$ | Production of c-di-GMP arising from the activated BphS* | c-di-GMP Signaling |
| R5 | $cdiGMP + PA2133 \xleftrightarrow{k_{5r},\ k_{5f}} cdiGMP\_PA2133$ | Degradation of c-di-GMP arising from the activated PA2133 | |
| R6 | $cdiGMP\_PA2133 \xrightarrow{k_6} PA2133$ | Degradation of c-di-GMP arising from the activated PA2133 | |
| R7 | $cdiGMP + FleQ \xleftrightarrow{k_{7r},\ k_{7f}} cdiGMP\_FleQ$ | Formation of the c-di-GMP dependent transcriptional complex | |
| R8 | $cdiGMP\_FleQ + cdiGMP \xleftrightarrow{k_{8r},\ k_{8f}} Di\_cdiGMP\_FleQ$ | Formation of the c-di-GMP dependent transcriptional complex | |
| R9 | $\emptyset \xrightarrow{k_9} cdiGMP$ | Production of c-di-GMP arising from the endogenous diguanylate cyclase in *Pseudomonas aeruginosa* | |
| R10 | $cdiGMP \xrightarrow{k_{10}} \emptyset$ | Degradation of c-di-GMP arising from the endogenous phosphodiesterase in *Pseudomonas aeruginosa* | |
| R11 | $PA10403 \xrightarrow{k_{11}} PA10403 + mRNA\_BphS$ | Transcription of BphS | BphS Expression |
| R12 | $mRNA\_BphS \xrightarrow{k_{12}} mRNA\_BphS + Di\_BphS$ | Translation of BphS | |
| R13 | $mRNA\_BphS \xrightarrow{\gamma_1} \emptyset$ | mRNA Degradation of BphS | |
| R14 | $Di\_BphS \xrightarrow{\gamma_2} \emptyset$ | Protein Degradation of Bphs dimer | |
| R15 | $Di\_BphS^* \xrightarrow{\gamma_2} \emptyset$ | Protein Degradation of Bphs* dimer | |
| R16 | $J23109 \xrightarrow{k_{13}} J23109 + mRNA\_PA2133$ | Transcription of PA2133 | PA2133 Expression |
| R17 | $mRNA\_PA2133 \xrightarrow{k_{14}} mRNA\_PA2133 + PA2133$ | Translation of PA2133 | |
| R18 | $mRNA\_PA2133 \xrightarrow{\gamma_1} \emptyset$ | mRNA Degradation of PA2133 | |
| R19 | $PA2133 \xrightarrow{\gamma_2} \emptyset$ | Protein Degradation of PA2133 | |
| R20 | $pCdrA + Di\_cdiGMP\_FleQ \xleftrightarrow{k_{15r},\ k_{15f}} pCdrA\_Di\_cdiGMP\_FleQ$ | C-di_GMP dependent transcriptional initiation | Q Expression |
| R21 | $pCdrA\_Di\_cdiGMP\_FleQ \xrightarrow{k_{16}} pCdrA\_Di\_cdiGMP\_FleQ + mRNA\_Q$ | Transcription of Q | |
| R22 | $mRNA\_Q \xrightarrow{k_{17}} mRNA\_Q + Q$ | Translation of Q | |
| R23 | $mRNA\_Q \xrightarrow{\gamma_1} \emptyset$ | mRNA Degradation of Q | |
| R24 | $Q \xrightarrow{\gamma_2} \emptyset$ | Protein Degradation of Q | |
| R25 | $pR' + Q \xleftrightarrow{k_{18r},\ k_{18f}} pR'\_Q$ | Q dependent transcriptional termination | LKD Expression |
| R26 | $pR'\_Q + Q \xleftrightarrow{k_{19r},\ k_{19f}} pR'\_Q\_Q$ | Q dependent transcriptional termination | |
| R27 | $pR'\_Q\_Q \xrightarrow{k_{20}} pR'\_Q\_Q + mRNA\_LKD$ | Transcription of LKD | |
| R28 | $mRNA\_LKD \xrightarrow{k_{21}} mRNA\_LKD + LKD$ | Translation of LKD | |
| R29 | $mRNA\_LKD \xrightarrow{\gamma_1} \emptyset$ | mRNA Degradation of LKD | |
| R30 | $LKD \xrightarrow{\gamma_2} \emptyset$ | Protein Degradation of LKD | |

#### Table S5. ODEs for the model reactions.

| # | Ordinary differential equation |
|---|---|
| Eq.1 | $\frac{d[Di\_BphS]}{dt} = -k_{1f}[photon][Di\_BphS]+k_{1r}[Di\_BphS\_photon] + k_3[Di\_BphS^*] + k_{10}[mRNA\_BphS] - \gamma_2[Di\_BphS]$ |
| Eq.2 | $\frac{d[Di\_BphS^*]}{dt} = k_{1f}[photon][Di\_BphS]-k_3[Di\_BphS^*] - \gamma_2[Di\_BphS^*]$ |
| Eq.3 | $\frac{d[cdiGMP]}{dt} = k_4[Di\_BphS^*]-k_{5f}[cdiGMP][PA2133]+k_{5r}[cdiGMP\_PA2133] - k_{7f}[cdiGMP][FleQ] + k_{7r}[cdiGMP\_FleQ] - k_{8f}[cdiGMP][cdiGMP\_FleQ] + k_{8r}[Di\_cdiGMP\_FleQ] + k_9 - k_{10}[cdiGMP]$ |
| Eq.4 | $\frac{d[PA2133]}{dt} = -k_{5f}[cdiGMP][PA2133] + k_{5r}[cdiGMP\_PA2133] + k_{14}[mRNA\_PA2133] + k_6[cdiGMP\_PA2133] - \gamma_2[PA2133]$ |
| Eq.5 | $\frac{d[cdiGMP\_PA2133]}{dt} = k_{5f}[cdiGMP][PA2133] - k_{5r}[cdiGMP\_PA2133] - k_6[cdiGMP\_PA2133]$ |
| Eq.6 | $\frac{d[cdiGMP\_FleQ]}{dt} = k_{7f}[cdiGMP][FleQ] - k_{7r}[cdiGMP\_FleQ] + k_{8r}[Di\_cdiGMP\_FleQ] - k_{8f}[cdiGMP][cdiGMP\_FleQ]$ |
| Eq.7 | $\frac{d[Di\_cdiGMP\_FleQ]}{dt} = k_{8f}[cdiGMP][cdiGMP\_FleQ] - k_{8r}[Di\_cdiGMP\_FleQ] - k_{15f}[pCdrA][Di\_cdiGMP\_FleQ] + k_{15r}[pCdrA\_Di\_cdiGMP\_FleQ]$ |
| Eq.8 | $\frac{d[mRNA\_BphS]}{dt} = k_{12}[PA1O4O3] - \gamma_1[mRNA\_BphS]$ |
| Eq.9 | $\frac{d[mRNA\_PA2133]}{dt} = k_{13}[J23109] - \gamma_1[mRNA\_PA2133]$ |
| Eq.10 | $\frac{d[pCdrA]}{dt} = -k_{15f}[pCdrA][Di\_cidiGMP\_FleQ] + k_{15r}[pCdrA\_Di\_cdiGMP\_FleQ]$ |
| Eq.11 | $\frac{d[pCdrA\_Di\_cdiGMP\_FleQ]}{dt} = k_{15f}[pCdrA][Di\_cdiGMP\_FleQ] - k_{15r}[pCdrA\_Di\_cdiGMP\_FleQ]$ |
| Eq.12 | $\frac{d[mRNA\_Q]}{dt} = k_{16}[pCdrA\_Di\_cdiGMP\_FleQ] - \gamma_1[mRNA\_Q]$ |
| Eq.13 | $\frac{d[Q]}{dt} = k_{17}[mRNA\_Q] - \gamma_2[Q] - k_{18f}[Q][pR'] - k_{19f}[Q][pR'\_Q] + k_{18r}[pR'\_Q] + k_{19r}[pR'\_Q\_Q]$ |
| Eq.14 | $\frac{d[pR']}{dt} = -k_{18f}[Q][pR'] + k_{18r}[pR'\_Q]$ |
| Eq.15 | $\frac{d[pR'\_Q]}{dt} = k_{18f}[Q][pR'] - k_{18r}[pR'\_Q] - k_{19f}[Q][pR'\_Q] + k_{19r}[pR'\_Q\_Q]$ |
| Eq.16 | $\frac{d[pR'\_Q\_Q]}{dt} = k_{19f}[Q][pR'\_Q] - k_{19r}[pR'\_Q\_Q]$ |
| Eq.17 | $\frac{d[mRNA\_LKD]}{dt} = k_{20}[pR'\_Q\_Q] - \gamma_1[mRNA\_LKD]$ |
| Eq.18 | $\frac{d[LKD]}{dt} = k_{21}[mRNA\_LKD] - \gamma_2[LKD]$ |

#### Table S6. Species and their initial conditions in the model.

| Species | Description | Unit | Initial Condition |
|---|---|---|---|
| $photon$ | Lighting power density | $\mu W cm^{-2}$ | 0.1 to 1000 |
| $Di\_BphS$ | Dimer of photon-activated diguanylate cyclase (BphS) at an inactivated state | $\mu M$ | 2 |
| $Di\_BphS^{*}$ | Dimer of photon-activated diguanylate cyclase ($BphS^{*}$) at an activated state | $\mu M$ | 0 |
| $cdiGMP$ | Second messenger cyclic diguanylate | $\mu M$ | 0 |
| $PA2133$ | Phosphodiesterase PA2133 for c-di-GMP degradation | $\mu M$ | 0.3 |
| $cdiGMP\_PA2133$ | Complex of the phosphodiesterase with c-di-GMP degradation | $\mu M$ | 0 |
| $FleQ$ | C-di-GMP dependent transcriptional factor FleQ | $\mu M$ | 1 |
| $cdiGMP\_FleQ$ | Complex of the FleQ with a c-di-GMP | $\mu M$ | 0 |
| $Di\_cdiGMP\_FleQ$ | Complex of the FleQ with two c-di-GMP | $\mu M$ | 0 |
| $PA1O4O3$ | Constitutive promoter PA1O4O3 | $\mu M$ | $1 \times 10^{-3}$ |
| $mRNA\_BphS$ | mRNA of BphS | $\mu M$ | 0 |
| $J23109$ | Constitutive promoter J23109 | $\mu M$ | $1 \times 10^{-3}$ |
| $mRNA\_PA2133$ | mRNA of PA2133 | $\mu M$ | 0 |
| $pCdrA$ | C-di-GMP dependent promoter pCdrA | $\mu M$ | 0.17 |
| $pCdrA\_Di\_cdiGMP\_FleQ$ | Transcriptional complex formed by the promoter pCdrA and the complex Di_cdiGMP_FleQ | $\mu M$ | 0 |
| $mRNA\_Q$ | mRNA of Q | $\mu M$ | 0 |
| $Q$ | Anti-terminator protein Q | $\mu M$ | 0 |
| $pR^{'}$ | Q dependent promoter pR‘ | $\mu M$ | $1 \times 10^{-3}$ |
| $pR^{'}\_Q$ | Transcriptional complex formed by the promoter pR’ and one Q | $\mu M$ | 0 |
| $pR^{'}\_Q\_Q$ | Transcriptional complex formed by the promoter pR‘and two Q | $\mu M$ | 0 |
| $mRNA\_LKD$ | mRNA of bacterial lysis genes | $\mu M$ | 0 |
| $LKD$ | Bacterial lysis proteins LKD | $\mu M$ | 0 |

#### Table S7. Kinetic constants obtained from the physiological ranges.

| Kinetic Constant | Description | Unit | Value | References/Notes |
|---|---|---|---|---|
| $k_{1f}$ | Binding constant arising from pseudo binding reaction of photon and BphS | $cm^2 \mu W^{-1} s^{-1}$ | | N.A. |
| $k_{1r}$ | Binding constant arising from pseudo dissociation reaction of the complex BphS_photon | $s^{-1}$ | 10 | N.A. |
| $k_2$ | Activation rate of the dimer of BphS in presence of NIR light | $s^{-1}$ | 80 | [14] |
| $k_3$ | Deactivation rate of the dimer of BphS in absence of NIR light | $s^{-1}$ | $1 \times 10^{-5}$ | [14] |
| $k_4$ | C-di-GMP production rate arising from activated BphS | $s^{-1}$ | 0.2 | N.A. |
| $k_{5f}$ | Binding constant of c-di-GMAP and PA2133 | $\mu M^{-1} s^{-1}$ | 100 | [29] |
| $k_{5r}$ | Dissociation rate of the complex formed by c-di-GMP and PA2133 | $s^{-1}$ | 1000 | N.A. |
| $k_6$ | C-di-GMP degradation rate arising from PA2133 | $s^{-1}$ | 12 | N.A. |
| $k_{7f}$ | Binding constant of c-di-GMAP and FleQ | $\mu M^{-1} s^{-1}$ | 100 | [29] |
| $k_{7r}$ | Dissociation rate of the complex formed by c-di-GMP and FleQ | $s^{-1}$ | 240 | [17] |
| $k_{8f}$ | Binding constant of c-di-GMAP and the complex formed by c-di-GMP and FleQ | $\mu M^{-1} s^{-1}$ | 100 | [29] |
| $k_{8r}$ | Dissociation rate of the complex formed by two c-di-GMP molecules and FleQ | $s^{-1}$ | 240 | [17] |
| $k_9$ | Production rate of c-di-GMP arising from the endogenous diguanylate cyclase in *Pseudomonas aeruginosa* | $\mu M s^{-1}$ | 0.2 | N.A. |
| $k_{10}$ | Degradation rate of c-di-GMP arising from the endogenous phosphodiesterase in *Pseudomonas aeruginosa* | $s^{-1}$ | $8 \times 10^{-2}$ | N.A. |
| $k_{11}$ | Transcriptional rate arising from the promoter of PA1O4O3 | $s^{-1}$ | $5 \times 10^{-2}$ | N.A. |
| $k_{12}$ | Translational rate arising from the mRNA of BphS | $s^{-1}$ | $5 \times 10^{-2}$ | N.A. |
| $\gamma_1$ | Degradation rate of mRNA | $s^{-1}$ | $5 \times 10^{-3}$ | [30] |
| $\gamma_2$ | Dilution rate of protein | $s^{-1}$ | $2.7 \times 10^{-4}$ | N.A. |
| $k_{13}$ | Transcriptional rate arising from the promoter of J23109 | $s^{-1}$ | $1.7 \times 10^{-3}$ | N.A. |
| $k_{14}$ | Translational rate arising from the mRNA of PA2133 | $s^{-1}$ | $1.2 \times 10^{-2}$ | N.A. |
| $k_{15f}$ | Binding constant of the promoter pCdrA and the transcriptional factor complex formed by two c-di-GMP molecule and FleQ | $\mu M^{-1} s^{-1}$ | 100 | [29] |
| $k_{15r}$ | Dissociation rate of the promoter pCdrA and the transcriptional factor complex formed by two c-di-GMP molecule and FleQ | $s^{-1}$ | 10 | N.A. |
| $k_{16}$ | Transcriptional rate arising from the c-di-GMP dependent promoter pCdrA | $s^{-1}$ | $6 \times 10^{-4}$ | N.A. |
| $k_{17}$ | Translational rate arising from the mRNA of Q | $s^{-1}$ | $3.5 \times 10^{-3}$ | N.A. |
| $k_{18f}$ | Binding constant of the promoter/terminator pR' and the anti-terminator Q | $\mu M^{-1} s^{-1}$ | 100 | [29] |
| $k_{18r}$ | Dissociation rate of the promoter/terminator pR' and the anti-terminator Q | $s^{-1}$ | 6 | [20] |
| $k_{19f}$ | Binding constant of the complex pR'_Q and the anti-terminator Q | $\mu M^{-1} s^{-1}$ | 100 | |
| $k_{19r}$ | Dissociation rate of the complex pR'_Q and the anti-terminator Q | $s^{-1}$ | 6 | [20] |
| $k_{20}$ | Transcriptional rate arising from the Q dependent promoter/terminator pR' | $s^{-1}$ | $2 \times 10^{-3}$ | N.A. |
| $k_{21}$ | Translational rate arising from the mRNA of LKD | $s^{-1}$ | $2 \times 10^{-2}$ | N.A. |

**Table S8. Dose schedule are set in the model.**

| Time (h) | Amount (micromole) | Rate (micromole/second) |
|---|---|---|
| 0 | 0 | 0 |
| 22 | 86.4 | 0.002 |
| 34 | 1728 | 0.04 |
| 46 | 86.4 | 0.002 |
| 58 | 1728 | 0.04 |
| 70 | 86.4 | 0.002 |
| 82 | 1728 | 0.04 |
| 94 | 86.4 | 0.002 |
| 106 | 1728 | 0.04 |
| 118 | 86.4 | 0.002 |
| 130 | 0 | 0 |

**Table S9. Experimental data used to determine model parameters.**

| Elements in the genetic circuit | | | | LPD range ($\mu W\cdot cm^{-2}$) | | |
|---|---|---|---|---|---|---|
| Pre-PA2133 | | Pre-Q | HlyE | Planktonic | Biofilm | Lysis |
| Promoter* | RBS1** | RBS2** | | | | |
| J23114 (0.1) | RBS010 (0.21) | - | - | 0-7.7 | >7.7 | - |
| J23105 (0.15) | RBS010 (0.21) | - | - | 0-7.7 | >7.7 | - |
| J23109 (0.04) | RBS010 (0.21) | - | - | 0-1.3 | >3.0 | - |
| J23109 (0.04) | RBS010 (0.21) | RBS017 (0.15) | + | 0-1.3 | 3.3-15.7 | >20 |
| J23109 (0.04) | RBS010 (0.21) | RBS 004 (0.28) | + | - | - | >2 |
| J23109 (0.04) | RBS010 (0.21) | RBS 016 (0.32) | + | - | - | >2 |
| J23109 (0.04) | RBS010 (0.21) | RBS 022 (0.17) | + | 0-1.3 | 3.3-15.7 | >20 |
| J23109 (0.04) | RBS010 (0.21) | RBS 036 (0.07) | - | 0-1.3 | 7.7-50.8 | >71.9 |

＊: Relative strength of promoter compared with BBa_J23100 (iGEM), and obtained from (https://parts.igem.org/Promoters/Catalog/Anderson)

＊＊: Relative strength of RBS library compared to BBa_B0034 (iGEM), and measured in this study.

#### Table S10. Relative strength and sequences of some RBS used in this study.

| RBS Name | Relative strength* | Base sequence | RBS Name | Relative strength | Base sequence |
|---|---|---|---|---|---|
| RBS002 | 0.4209 | AAACTGGCCCAA | RBS041 | 0.0309 | CGAGACCAGAAG |
| RBS004 | 0.2834 | AACGGGGATGAA | RBS042 | 0.0376 | CCAGAACAGAAA |
| RBS005 | 0.0304 | AAGTCGGCAAAA | RBS045 | 0.0088 | CGCAGTGTCGGC |
| RBS006 | 0.0640 | AAGCTCATCGAA | RBS046 | 0.4628 | GAAGCAAAGGGG |
| RBS007 | 0.0164 | AATCCTACGAAA | RBS047 | 0.0169 | TCGTGTGCCGTT |
| RBS008 | 0.0292 | AAACCAATCAAA | RBS048 | 0.0463 | ATCGCAAGGAAA |
| RBS009 | 0.0022 | AAGGACAAGAAA | RBS049 | 0.0381 | GAGTAGTGCAAG |
| RBS010 | 0.2119 | AAATTTGGGAAA | RBS050 | 0.0154 | TGTTCGCTGAGT |
| RBS011 | 0.9628 | AAAAAAGGGGAA | RBS051 | 0.0651 | GTGGAACCAAGA |
| RBS012 | 0.1282 | AAGGGCAGGCAA | **RBS052****| **1.0000** | **AAAGAGGAGAAA** |
| RBS013 | 0.0491 | GAAGACTAGAGC | RBS053 | 0.0080 | GTTGCCACGACA |
| RBS014 | 0.1862 | GGAGAATAGAGT | RBS054 | 0.0319 | TGGTTTGAGGGA |
| RBS015 | 0.0634 | AATGTGGCGGAA | RBS055 | 0.0112 | CTCAAACATCGC |
| RBS016 | 0.3249 | AAGGAGGGGGAA | RBS056 | 0.0611 | CAACGCTTGAGG |
| RBS017 | 0.1466 | AACGCGGTGCAA | RBS057 | 0.1378 | ATTGGCTTTGGG |
| RBS018 | 0.6692 | AAAGTGGAGTAA | RBS058 | 0.0114 | GTCTAGTATCCA |
| RBS019 | 0.1012 | AAAGGGGCGAAA | RBS059 | 0.0204 | ATGTGTCTGGTC |
| RBS020 | 0.3734 | AACGGGGTGTAA | RBS060 | 0.0481 | GGGAGAATGCTT |
| RBS021 | 0.0706 | AAGGCGGTCCAA | RBS062 | 0.0073 | CACCCTATAGGG |
| RBS022 | 0.1678 | AATGAGGGTGAA | RBS063 | 0.0094 | GCTTCGTCGGGC |
| RBS023 | 0.1192 | AAGCTGGCTGAA | RBS064 | 0.0095 | CACTCCGTCCCA |
| RBS024 | 0.1261 | AATGAGGCCGAA | RBS065 | 0.0414 | CCCGGACTAGAC |
| RBS025 | 0.1301 | AAATGGGCTAA | RBS066 | 0.0192 | CATGTACCACGT |
| RBS026 | 0.0548 | AACACGGCTGAA | RBS067 | 0.1294 | AAATTTGCAGGA |
| RBS027 | 0.0307 | AATCCGGACGAA | RBS068 | 0.0279 | TTCAAGGTGCGG |
| RBS028 | 0.0406 | AATCGGGTCGAA | RBS070 | 0.1025 | CGCAATCTATCG |
| RBS029 | 0.0332 | AAATAGTACCAA | RBS071 | 0.0097 | CTCGAATTGCAT |
| RBS030 | 0.0210 | AACCCGTACCAA | RBS073 | 0.0090 | GCGTTGTGCCTG |
| RBS031 | 0.0230 | AAGCGTTAAAAA | RBS074 | 0.0072 | CATATAACGCCT |
| RBS032 | 0.0194 | AAACCCCGCTAA | RBS075 | 0.0723 | CAAACCTATTGT |
| RBS033 | 0.0549 | AAGAACGTGTAA | RBS076 | 0.0119 | GGTCTCGGCACA |
| RBS035 | 0.0129 | AAGCCGGATCAA | RBS077 | 0.0726 | GGATTGGAACGG |
| RBS036 | 0.0755 | AAGGCGATCAAA | RBS078 | 0.0072 | TCGAGTTTTACG |
| RBS037 | 0.0309 | CCAGATTAGAAC | RBS079 | 0.0085 | TGAGCGCATACC |
| RBS038 | 0.1529 | AAAGAAGAGAAT | RBS080 | 0.0070 | AGCCTCCATCGT |
| RBS040 | 0.0973 | GAAGAAGAGACT | RBS081 | 0.0114 | TAACTTCCCAA |

*: Relative strength compared to BBa_B0034 (iGEM).

**: The base sequence of RBS052 is exactly the same as that of B0034.

#### Table S11. Bacterial Strains and plasmids used in this study.

| | Relevant characteristics | Source |
|---|---|---|
| ***P. aeruginosa* strains** | | |
| PAO1 | Wild-type strain. None resistance | J.D. Shrout |
| pR'-mScarlet | PAO1, pR'-B0034-*mScarlet*-Tn7. None resistance | This study |
| pR'-tR'-mScarlet | PAO1, pR'-tR'-B0034-*mScarlet*-Tn7. None resistance | This study |
| pR'-tR'-mScarlet/Q-pJN105 | pR'-tR'-mScarlet containing plasmid *Q*-pJN105. Gm$^{r}$ | This study |
| PAO1-BphS | PAO1, P*A1*/*O4*/*O3*-RBSII-*bphS*-CTX2. None resistance | This study |
| PAO1-BphS-J23114-PA2133 | PAO1-BphS, J23114-RBS010-*PA2133*-Tn7. Gm$^{r}$ | This study |
| PAO1-BphS-J23105-PA2133 | PAO1-BphS, J23105-RBS010-*PA2133*-Tn7. Gm$^{r}$ | This study |
| PAO1-BphS-J23109-PA2133 | PAO1-BphS, J23109-RBS010-*PA2133*-Tn7. Gm$^{r}$ | This study |
| PAO1-BphS-LKD | PAO1-BphS, pR'-tR'-B0034-*LKD*-J23019-RBS010-*PA2133*-Tn7, None resistance | This study |
| PcdrA-GFP | PAO1-BphS-LKD, P*cdrA*-*gfp*(mut3)-PUCP20. Gm$^{r}$ | This study |
| RBS004-Q | PAO1-BphS-LKD, P*cdrA*-*gfp*(mut3)-PcdrA-RBS004-*Q*(ASV)-pUCP20. Gm$^{r}$ | This study |
| RBS018-Q | PAO1-BphS-LKD, P*cdrA*-*gfp*(mut3)-PcdrA-RBS018-*Q*(ASV)-pUCP20. Gm$^{r}$ | This study |
| PAO1-LKD | PAO1, pR'-tR'-B0034-*LKD*-J23019-RBS010-*PA2133*-Tn7, None resistance | This study |
| PAO1-BphS-LKD-GFP | PAO1-LKD, P*A1*/*O4*/*O3*-RBSII-*bphS*-J23102-B0034-*sfGFP*-CTX2. None resistance | This study |
| RecA | *vfr , and recA* double gene knockouts in PAO1-BphS-LKD-GFP. | This study |
| RecA-H036 | RecA, J23118-RBSII-*hlyE*-T0T1-PcdrA-RBS036-*Q*(ASV) -pUCP20. Gm$^{r}$ | This study |
| ExoST | *vfr ,exoS and exoT* triple gene knockouts in PAO1-BphS-LKD-GFP. | This study |
| H017 | ExoST, J23118-RBSII-*hlyE*-T0T1-P*cdrA*-RBS017-*Q*(ASV) -pUCP20. Gm$^{r}$ | This study |
| Q017 | ExoST, P*cdrA*-RBS017-*Q*(ASV) -pUCP20. Gm$^{r}$ | This study |
| M023 | ExoST, J23118-RBSII-*mScarlet*-T0T1-PcdrA-RBS023-*Q*(ASV) -pUCP20. Gm$^{r}$ | This study |
| ***E.coli* strains** | | |
| Top 10 | F−, mcrA, (mrr, hsdRMS–mcrBC), 80lacZ M15 lacX74, recA1, araD139, (ara–leu)7697, galU, galK, rpsL(StrR), endA1, nupG. None resistance | Invitrogen |
| **Plasmids** | | |
| pJN105 | pJN105 araC-PBAD cassette cloned in pBBR1MCS-5. Gm$^{r}$ | This study |
| pUC18T-mini-Tn7T-Gm | Mini-Tn7 transposon vector. Gm$^{r}$ | H.P.Schweizer |
| pTNS2 | a helper plasmid for mini-Tn7 site-specific transposition system. Ap$^{r}$ | H.P.Schweizer |
| pFLP2 | FRT cassette vector for Flp recombinase. Ap$^{r}$ | H.P.Schweizer |
| pEX18Gm | Gene replacement vector with MCS from pUC18. Gm$^{r}$ | H.P.Schweizer |
| miniCTX2 | mini-CTX2 transposon vector. Tet$^{r}$ | H.P.Schweizer |
| pEX18Gm-*vfr* | pEX18Gm-derived allelic-exchange vector for *vfr*. Gm$^{r}$ | This study |
| pEX18Gm-*recA* | pEX18Gm-derived allelic-exchange vector for *recA*. Gm$^{r}$ | This study |
| pEX18Gm-*exoS* | pEX18Gm-derived allelic-exchange vector for *exoS*. Gm$^{r}$ | This study |
| pEX18Gm-*exoT* | pEX18Gm-derived allelic-exchange vector for *exoT*. Gm$^{r}$ | This study |
| PA1/O4/O3-RBSII-*bphS*-CTX2 | Plasmid used for chromosomal insertion of *bphS* at attB site. Tet$^{r}$ | This study |
| J23114-RBS010-*PA2133*-Tn7 | Plasmid used for tuning expression level of *PA2133* with promoter J23114 and RBS010 | This study |
| J23105-RBS010-*PA2133*-Tn7 | Plasmid used for tuning expression level of *PA2133* with promoter J23105 and RBS010 | This study |
| J23109-RBS010-*PA2133*-Tn7 | Plasmid used for tuning expression level of *PA2133* with promoter J23109 and RBS010 | This study |
| pR'-tR'-B0034-*LKD*-J23019-RBS010-*PA2133*-Tn7 | Plasmid used for chromosomal insertion of *LKD* and *PA2133* at attTn7 site. Gm$^{r}$ | This study |
| P*cdrA*-*gfp*(mut3)-pUCP20 | c-di-GMP reporter plasmid. Gm$^{r}$ | This study |
| P*cdrA*-*gfp*(mut3)-PcdrA-RBS018 - *Q*(ASV)-pUCP20 | Plasmid used for detection of intracellular c-di-GMP level together with *Q* downstream of RBS018. Gm$^{r}$ | This study |
| P*cdrA*-*gfp*(mut3)-P*cdrA*-RBS004-*Q*(ASV)-pUCP20 | Plasmid used for detection of intracellular c-di-GMP level together with *Q* downstream of RBS004. Gm$^{r}$ | This study |
| PA1/O4/O3-RBSII-*bphS*-J23102-B0034-*sfGFP*-CTX2 | Plasmid used for chromosomal insertion of *bphS* and *sfGFP* at attB site. Tet$^{r}$ | This study |
| J23118-RBSII-*mScarlet*-T0T1-P*cdrA*-RBS017-*Q*(ASV) - PUCP20 | Plasmid used for expression of *mScarlet* and *Q*. Gm$^{r}$ | This study |
| J23118-RBSII-*hlyE*-T0T1-P*cdrA*-RBS017-*Q*(ASV) -PUCP20 | Plasmid used for expression of *hlyE* and *Q*. Gm$^{r}$ | This study |
| J23118-RBSII-*hlyE*-T0T1-P*cdrA*-RBS036-*Q*(ASV) -PUCP20. | Plasmid used for expression of *hlyE* and *Q*. Gm$^{r}$ | This study |
| P*cdrA*-RBS017-*Q*(ASV) -PUCP20 | Plasmid used for expression of *Q* under control of P*cdrA* ptomoter and RBS017. Gm$^{r}$ | This study |
| J23105-B0034-*sfGFP*-T0T1-J23102-*cyOFP*-pUCP20 | Plasmid used as a template to construct a RBS mutant library in *P. aeruginosa.* Gm$^{r}$ | This study |
| pR'-B0034-*mScarlet*-Tn7 | Plasmid used for chromosomal insertion of pR'-B0034-*mScarlet* fragment at attTn7 site. Gm$^{r}$ | This study |
| pR'-tR'-B0034-*mScarlet*-Tn7 | Plasmid used for chromosomal insertion of pR'-tR'-B0034-*mScarlet* at attTn7 site. Gm$^{r}$ | This study |
| *Q*-pJN105 | pJN105 inducible vector with fragment of *Q*. Gm$^{r}$ | This study |

**Movie S1. Time-lapse bright field images of H017 illuminated with Low-LPD NIR at 60× magnification.**

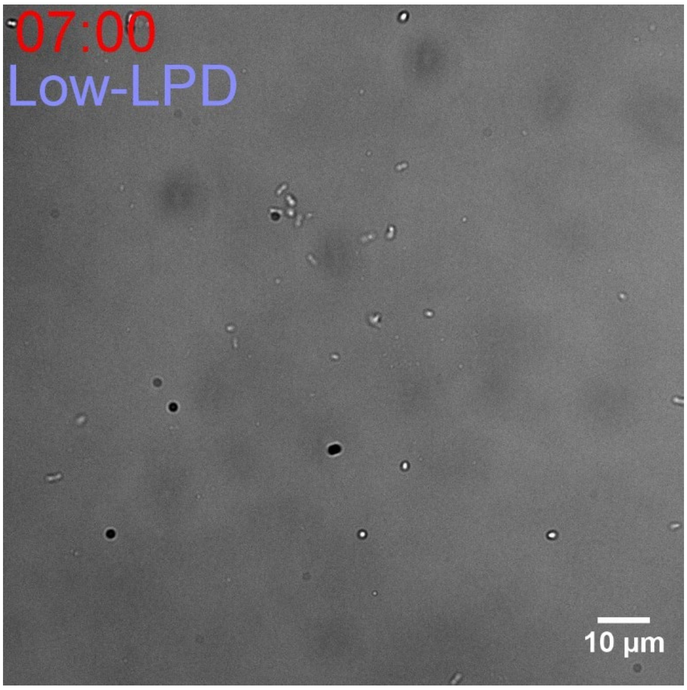

**Movie S2. Time-lapse bright field images of H017 illuminated with Medium-LPD NIR at 60× magnification.**

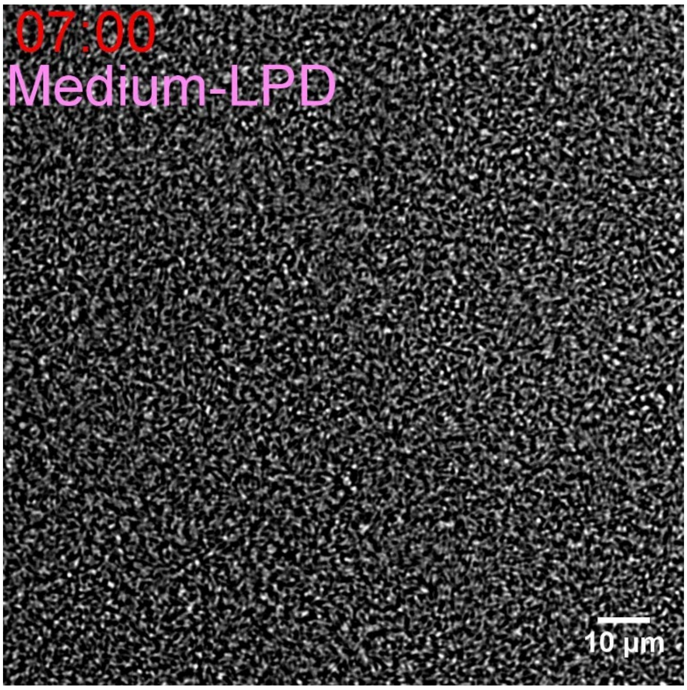

**Movie S3. Time-lapse bright field images of H017 illuminated with High-LPD NIR at 60× magnification.**

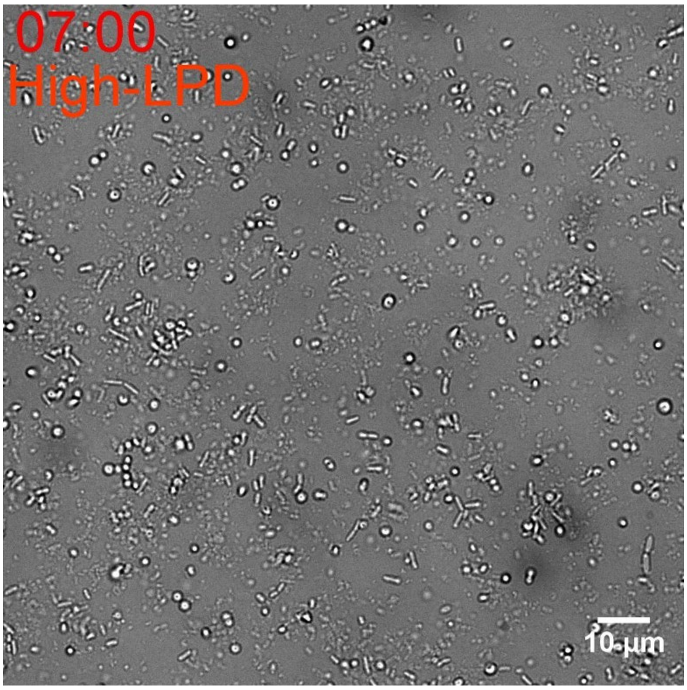

**Movie S4. Time-lapse fluorescence images of H017 illuminated with periodical Medium-LPD NIR and High-LPD NIR for 5 cycles at 60 × magnification.**

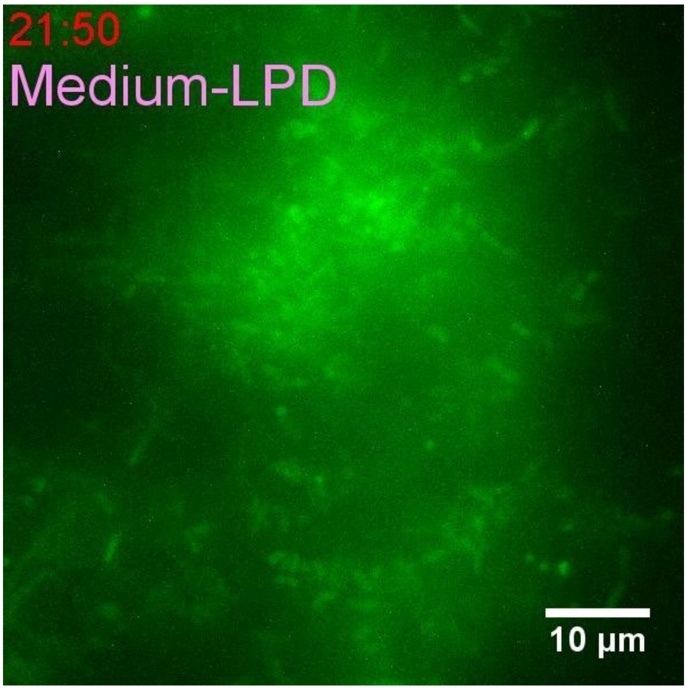

**Movie S5. H017 and live A549 cells co-cultured on a microfluidic device and illuminated with Medium-LPD NIR first and High-LPD NIR later at 60 × magnification.**

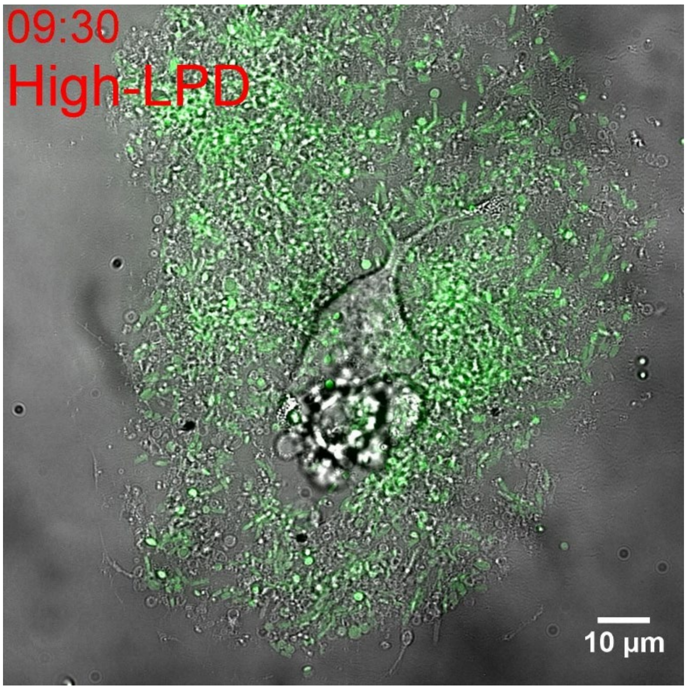

**Movie S6. H017 and live A549 cells co-cultured on a microfluidic device illuminated with High-LPD NIR at 60× magnification.**

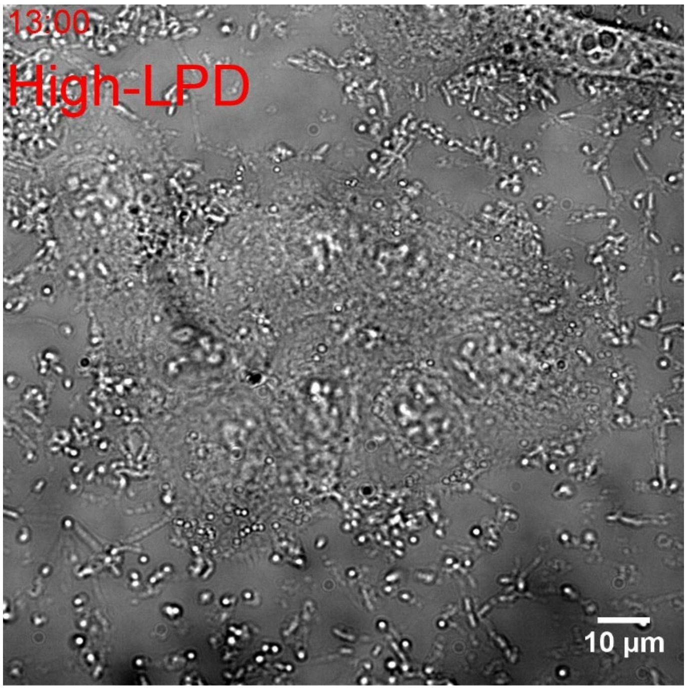

**Movie S7. ExoST and live A549 cells co-cultured on a microfluidic device and illuminated with Medium-LPD NIR at 60 × magnification.**

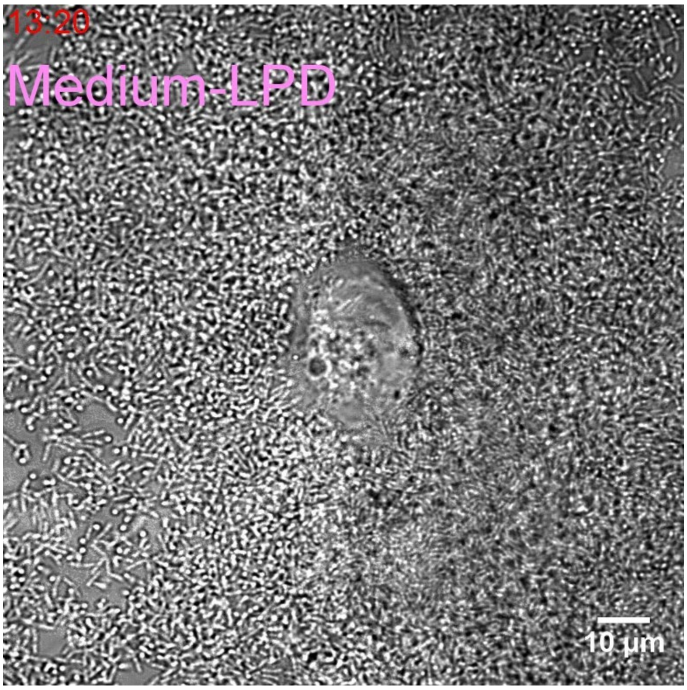
